# Functional segregation of body-brain signals in the area postrema

**DOI:** 10.64898/2026.06.15.732473

**Authors:** Alejandro López-Cruz, Natalie S. Figueredo Burgos, Anna M. Hakimi, Kathryn Xie, Mo Mao, Mahekdeep Kaur, Antoinette Spina, Katie Choi, Longhui Qiu, Teresa E. Lever, Fiona M. Gribble, Frank Reimann, Martin G. Myers, Alice E. Adriaenssens, Zachary A. Knight

**Author notes:** Corresponding author (ZAK).

## Abstract

Nausea arises from activation of specialized neurons in the area postrema (AP)^1–8^. The AP also mediates much of the satiety produced by GLP1R agonists^9–12^, suggesting a broader role in non-aversive physiology, yet the functions of AP cell types are not well understood. Here, we have used optical recordings in behaving mice to systematically define the natural regulation of an array of AP neurons, including the cell types that are principal targets of widely-used weight loss drugs. We discover that neurons expressing GFRAL, the receptor for the sickness-related hormone GDF15, are unexpectedly activated when mice consume food rich in fat. This fat-specific GFRAL neuron activation is required for fat satiation but does not involve GDF15 or canonical gut-brain pathways. Instead, “anti-nausea” neurons expressing GIPR, which directly inhibit GFRAL neurons, are selectively activated by sugar, enabling macronutrient-specific gating of GFRAL responses. In addition, we show that CALCR neurons link intestinal hyperosmolality to the suppression of feeding, whereas PRLHR neurons respond to changes in blood volume and pressure. These findings reveal a broad role for AP cell types in sensing and responding to physiologic signals unrelated to nausea. They also reveal that GFRAL and GIPR neurons, which are key targets of the weight-loss drug tirzepatide, have a natural function in sensing ingestion of fat and sugar, respectively.

## Introduction

The area postrema (AP) is a small structure in the caudal brainstem that is critical for nausea. The AP is situated outside the blood-brain barrier, and AP neurons express receptors for a variety of blood-borne substances that can induce sickness. Stimulation of AP cell types induces nausea-like aversive responses in mice^7,8^, whereas AP lesions or silencing prevent the sickness induced by emetics in cats^1,2,13^, dogs^14–16^, rodents^17–19^, and humans^20^. Thus, the AP is thought to be specialized for detecting noxious signals in the blood and then transforming these signals into sickness responses.

Nevertheless, the growing realization that the AP is the critical target of weight loss drugs based on glucagon-like peptide-1 (GLP1) and other peptides^9–12^, which can suppress appetite independent of nausea^21–23^, has stimulated interest in the role of this structure in non-aversive physiology. However, a fundamental challenge has been the lack of information about the natural regulation of AP circuits. To date, there have been no recordings of the activity of any AP cell type, in any organism, during behavior. This is due to the location of the AP in the extreme caudal brainstem, which creates unique challenges for performing neural recordings in awake animals. Thus, it remains generally unknown whether and how AP cell types respond to a broad range of physiologic signals unrelated to sickness.

The AP contains a variety of molecularly distinct cell types^7,24^. One attractive, but untested model is that these genetically-defined cells respond to distinct signals during behavior and, in turn, control different aspects of physiology. Here, we test this idea by using optical recordings in behaving mice to define the physiologic function of a panel of AP cell types, including the cells that are the principal targets of each major class of peptide hormone drugs.

## Results

### AP cell types show distinct response profiles to a panel of physiologic stimuli

The AP contains seven molecularly-distinct cell types^7,24^, four of which are glutamatergic and three of which are GABAergic (Fig. 1a). To investigate the natural function of these cells, we assembled Cre drivers that provide selective genetic access to five of these seven cell types (Fig. 1a). Of note, these five cell types encompass, collectively, all the AP cells that express receptors for the three major classes of peptide weight loss drugs (glucagon-like peptide-1 receptor, GLP1R; amylin receptor, AMYR; and gastric inhibitory polypeptide receptor, GIPR; Fig. 1b,c).

**Figure 1.**
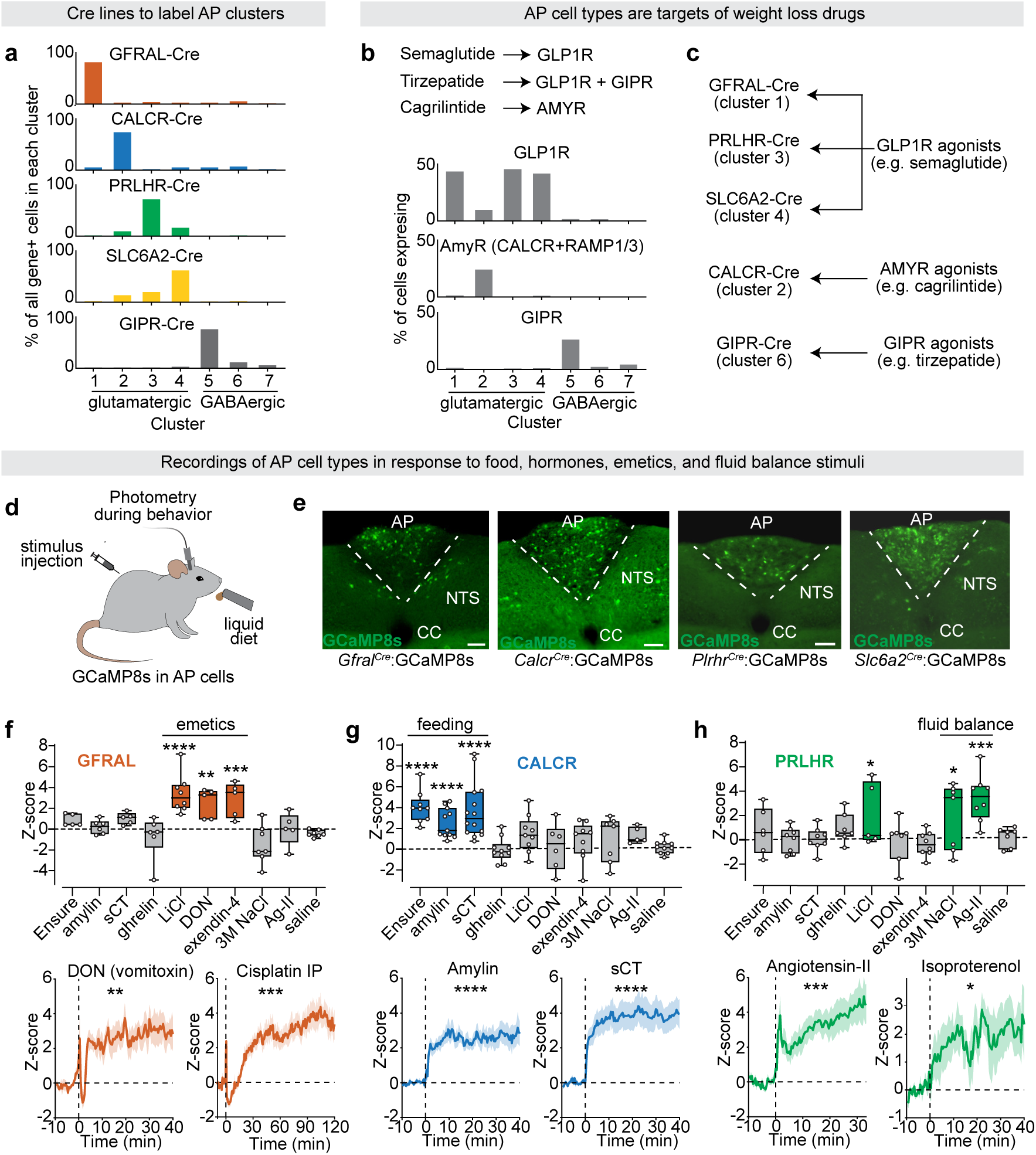
Area postrema (AP) cell types are tuned to different physiologic stimuli. **a,** Expression of each Cre line gene (*Gfral, Calcr, Prlhr, Slc6a2, or Gipr*) in each AP cluster, analyzed from Zhang et al. Cluster numbers were re-arranged for presentation clarity. Plot shows the percent of all Cre gene positive cells that are in each cluster. **b,** Percent of cells in each cluster expressing the indicated receptor. AmyR positive cells express both *Calcr* and either *Ramp1* or *Ramp3.* **c,** Schematic showing how the five Cre drivers encompass, collectively, all of the AP cells that express receptors for the three major classes of peptide weight loss drugs. **d,** Schematic of fiber photometry recordings of AP neurons in response to IP injections and Ensure ingestion. **e,** GCaMP8s expression in the four glutamatergic AP cell types. NTS: nucleus tractus solitarius. CC: central canal. Scale bar: 100um. **f,** Top, Boxplot showing average photometry z-score (0-40 min) after IP injection or Ensure access for GFRAL neurons. Bottom, GFRAL neuron response aligned to IP injection of indicated emetic. **g,** Top, Boxplot showing average photometry z-score (0-40 min) after IP injection or Ensure access for CALCR neurons. Bottom, CALCR neuron response aligned to IP injection of indicated hormones. **h,** Top, Boxplot showing average photometry z-score (0-40 min) after IP injection or Ensure access for PRLHR neurons. Bottom, PRLHR neuron response aligned to IP injection of indicated fluid balance stimuli. Colored boxes in f-h are statistically significant. Statistical comparisons are relative to the baseline prior to injection/ingestion. *P<0.05, **P<0.01, ***P<0.001, ****P<0.0001. Data are mean ± SEM.

We first investigated the function of the four glutamatergic cell types, as these include all the projection neurons of the AP^7^. We targeted GCaMP8s to the AP in each Cre line and implanted an angled optical fiber above the AP for fiber photometry recordings (Fig. 1d,e). We then challenged mice with injections of a panel of hormones, emetic agents, and hemodynamic stimuli or allowed them to consume the liquid diet Ensure. We found that neurons expressing GDNF family receptor alpha-like (GFRAL, cluster 1), the receptor for growth differentiation factor 15 (GDF-15), were consistently activated by emetics (LiCl, vomitoxin, high-dose Exendin-4, and cisplatin) but did not respond to signals associated with energy or fluid balance (Fig. 1f, Extended Data Fig 1a). Neurons expressing calcitonin receptor (CALCR, cluster 2) responded robustly to the nutritionally-regulated hormones calcitonin and amylin, consistent with their expression of the amylin receptor, and were also activated by Ensure ingestion, but showed no responses to emetics or hemodynamic signals (Fig. 1g, Extended Data Fig 1b). Prolactin-releasing hormone receptor (PRLHR) neurons (cluster 3), on the other hand, were selectively activated by a broad range of signals associated with fluid balance, including injection of Angiotensin-II (Ag-II), which is released in response to hypovolemia, as well as by injection of hypertonic solutions of NaCl and LiCl, which produce dehydration (Fig. 1h, Extended Data Fig 1c). We confirmed that PRLHR neurons sense physiologic changes in fluid balance by showing that they are activated by hypovolemia in an Ag-II-dependent manner (Extended Data Fig 1d,e) as well as by hypotension (Fig. 1h, isoproterenol). Finally, we found that neurons expressing norepinephrine transporter (SLC6A2, cluster 4) were strongly activated by exendin-4, consistent with high expression of GLP1R in this cluster^7^, but responded inconsistently to other stimuli (Extended Data Fig 1f,g). Taken together, these data reveal that molecularly-defined AP cell types are tuned to stimuli associated with different aspects of physiology (food ingestion, sickness, and hemodynamic state).

### GFRAL neurons are rapidly activated by emetics through GDF-15 independent mechanisms

GFRAL neurons were the only cell type in our screen that responded consistently to emetics (Fig. 1), which is the function traditionally ascribed to the AP^1–8^. We therefore investigated first how GFRAL neurons detect sickness-associated signals and whether they are engaged in other physiologic contexts.

GFRAL is the receptor for the hormone GDF15^25–27^, which is produced by peripheral tissues in response to diverse stimuli that promote nausea and sickness^25,28–39^, and we confirmed that intraperitoneal (IP) administration of GDF-15 activates GFRAL neurons as measured by photometry (2.38 ± 0.41 for GDF-15 vs. 0.09 ± 0.43 for vehicle, p=0.0027; Fig. 2a). GDF-15 synthesis and secretion occurs over hours^34,40–42^, but the activation of GFRAL neurons by emetics in our screen was much faster (Fig. 2b-d, Extended Data Fig 2a,b). This activation occurred in two phases: an initial spike in GFRAL activity at the moment of injection, which may reflect injection stress (Fig. 2b,c, zoomed rectangle), followed by a second phase response that occurred with a variable delay of a few minutes (e.g. LiCl: 3.2 ± 0.5 min; DON: 1.5 ± 0.3 min; cisplatin: 13.8 ± 1.0 min, exendin-4: 6.0 ± 3.2 min; Fig. 2b-d, Extended Data Fig 2a,b).

**Figure 2.**
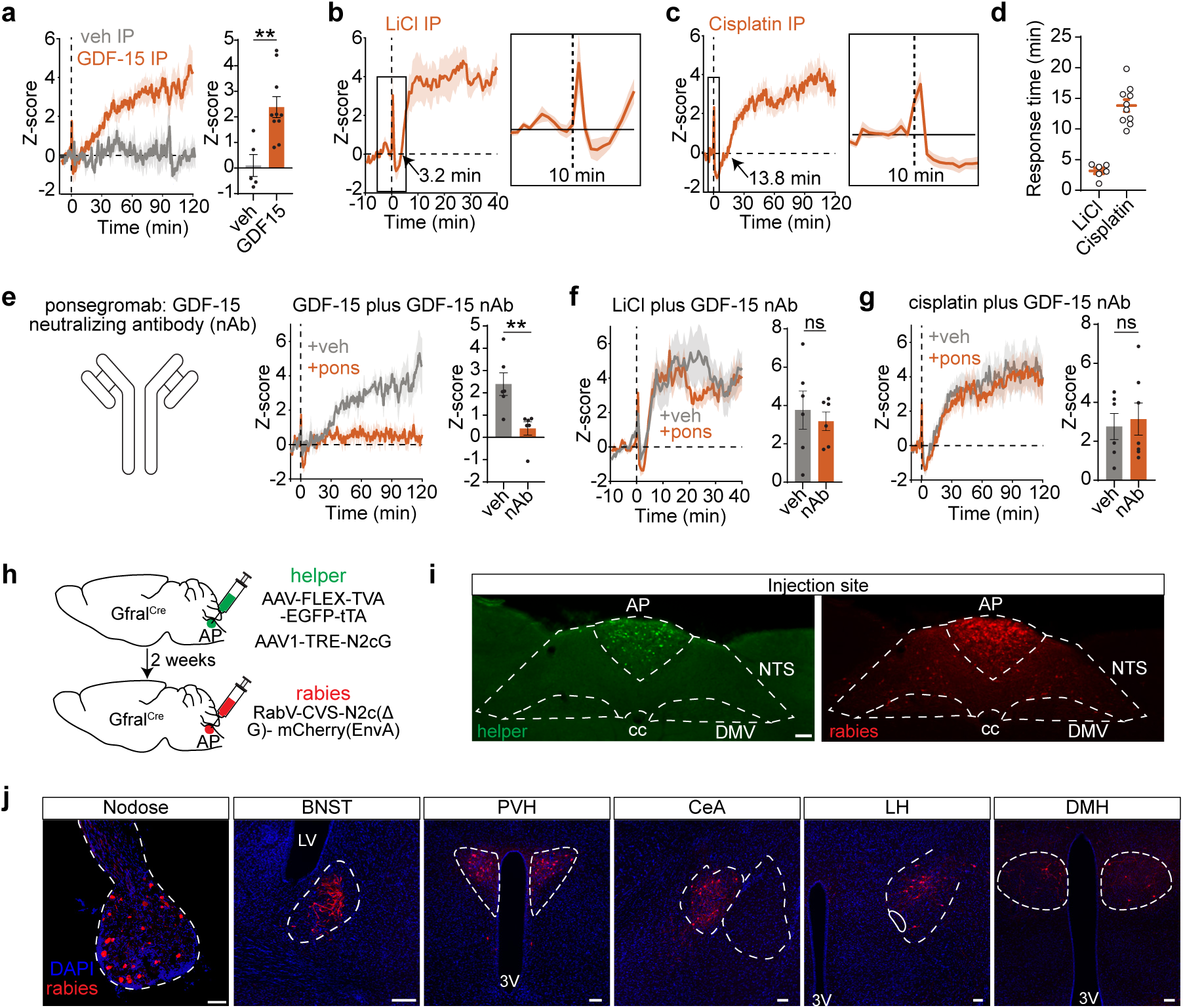
GFRAL neurons respond rapidly to emetics, independently of GDF-15. **a,** GFRAL neuron response aligned to IP injection of the GFRAL ligand, GDF-15 (positive control). **b,** GFRAL neuron responses aligned to IP injection of the emetic LiCl. GFRAL activation occurred in two phases: an initial spike in GFRAL activity at the moment of injection, followed by a second sustained phase response that occurred with a variable delay of a few minutes. Zoom box shows initial activation, 5 min before and after IP injection. Arrow indicates average response time of second sustained phase of activation. **c,** Response to IP cisplatin. **d,** Response time for second activation phase in individual animals to LiCl and cisplatin injection. **e,** GFRAL neuron responses to GDF-15 in animals pre-treated with vehicle or ponsegromab (GDF-15 neutralizing antibody). **f,** Left, GFRAL neuron responses to LiCl in animals pre-treated with vehicle or ponsegromab. Right. z-scores. **g,** Left, GFRAL neuron responses to cisplatin in animals pre-treated with vehicle or ponsegromab. Right. z-scores. **h,** Schematic for monosynaptic rabies tracing experiment. Animals were injected directly in the area postrema with two helper viruses encoding rabies TVA and G protein. Two weeks later CVS-N2c(ΔG) rabies virus was injected into the AP in a second surgery. Animals were perfused and tissue was processed 7 days later. **i,** Left, expression of helper viruses (green) is restricted to the AP. DMV: dorsal motor vagus.Right. Rabies virus expression in AP and NTS (red); scale bar: 100μm. **j,** Monosynaptic inputs onto GFRAL neurons include nodose ganglion, bed nucleus of the stria terminalis (BNST), paraventricular hypothalamus (PVH), central amygdala (CeA), lateral hypothalamus (LH), and dorsomedial hypothalamus (DMH). LV: lateral ventricle. 3V: third ventricle. scale bar: 100μm. ns, not significant, **P<0.01, ****P<0.0001. Data are mean ± SEM.

To test whether this second phase requires GDF-15, we pretreated animals with ponsegromab, a high-affinity neutralizing antibody against GDF-15^43,44^. We found that ponsegromab abolished GFRAL neuron responses to injection of exogenous GDF-15 (Fig. 2e), confirming its efficacy, but otherwise had no effect on the response to either LiCl or cisplatin (which is known to induce GDF-15 secretion^32,39^) (3.14 ± 0.83z for cisplatin plus ponsegromab vs. 2.75 ± 0.67z for cisplatin plus vehicle, p=0.84; Fig. 2f,g). This indicates that emetics can rapidly activate GFRAL neurons through GDF-15-independent mechanisms.

The AP is thought to respond primarily to humoral signals^4–6^, but the fast responses we observe in GFRAL neurons suggests a possible role for synaptic inputs.. To identify the source of synaptic inputs, we performed monosynaptic rabies tracing from these cells. Helper viruses encoding TVA and rabies glycoprotein were injected directly into the AP of GFRAL-Cre mice, followed two weeks later by injection of EnvA-pseudotyped, glycoprotein-deleted rabies virus (Fig. 2h). We confirmed that this procedure selectively infected cells in the AP without spread into the adjacent NTS (Fig. 2i). We observed labeled presynaptic neurons in the nodose ganglia, indicating that GFRAL neurons receive input from the periphery through the vagus nerve, as well as labelled cells in several forebrain structures associated with stress and energy balance, including the paraventricular hypothalamus (PVH), bed nucleus of the stria terminalis (BNST), central amygdala (CeA), dorsomedial hypothalamus (DMH), and lateral hypothalamus (LH) (Fig. 2j). This provides a pathway for rapid non-humoral activation of GFRAL neurons by aversive or other stimuli.

### GFRAL neurons are activated by ingestion of fat

Although GFRAL neurons are strongly associated with aversion, we wondered whether there might be a context in which these cells are activated during normal ingestion. To investigate this, we fasted mice overnight and then recorded GFRAL neuron responses during self-paced consumption of different liquid diets (Fig. 3a-c). To our surprise, we found that these cells were dramatically activated when hungry mice consumed a pure fat solution (Intralipid 10%). This activation ramped over the one-hour recording and tracked cumulative licks (R^2^ = 0.38, p < 0.001) but not instantaneous lick rate or recent licking history (Fig. 3d), suggesting GFRAL neurons track total fat consumed rather than ingestion rate. Varying the duration of Intralipid access confirmed that responses scaled linearly with total fat consumed (R^2^ = 0.32, p= 0.017; Extended Data Fig 3a-c). In contrast, we observed no activation when mice consumed a pure sugar solution (24% glucose) and only weak responses to a mixed diet that is 25% fat (0.85 ± 0.26z for Ensure, 3.54 ± 0.74z for Intralipid, 0.10 ± 0.48z for glucose, p=0.002, Fig. 3b-c), despite no differences in total consumption (3.28 ± 0.28 kcal for Ensure vs. 2.90 ± 0.14 kcal for Intralipid, p=0.35; Fig.3c). Thus, GFRAL neurons are strongly and selectively activated by consumption of fat.

**Figure 3.**
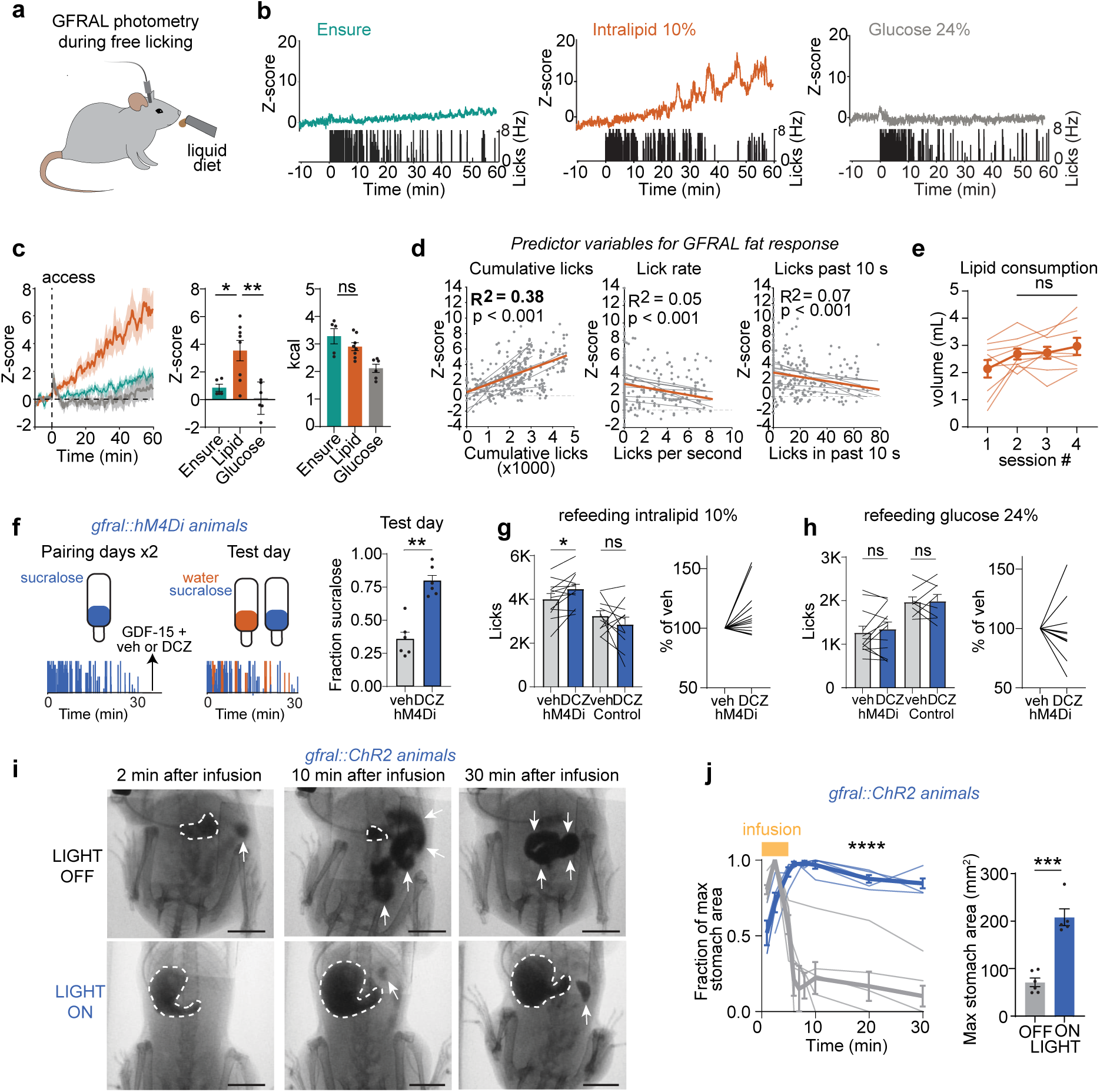
GFRAL neurons are activated by dietary fat, and pace ingestion and suppress gastric emptying. **a,** Schematic depicting fiber photometry recordings during free licking. **b,** Example trace of calcium dynamics of GFRAL neurons during self-paced consumption of Ensure, intralipid 10%, or glucose 24%. The lick rate is shown below. **c,** Left. GFRAL neuron activity aligned to access to different macronutrients. Right. z-score and kcal consumed (0-60 min). **d,** Regression analysis for predictor variables (cumulative licks, lick rate, licks in past 10 sec) vs z-score during intralipid licking. **e,** Volume of intralipid consumed across different 60 min sessions on different days. Sessions were separated by at least two days. **f,** Left, schematic for conditioned taste aversion to GDF-15. Sucralose was paired with GDF-15 and either vehicle or DCZ injection. Animals then underwent two bottle choice test. Right, Fraction time spent licking sucralose (vs. water) during two bottle choice. **g,** Left, intralipid consumption (during 60 min) after either vehicle or DCZ injection in *Gfral::hM4Di* animals or control littermates. Right, change in intralipid consumption after DCZ relative to vehicle in *Gfral::hM4Di* animals. **h,** Left, glucose consumption (during 60 min) after either vehicle or DCZ injection in *Gfral::hM4Di* or control littermates. Right, change in glucose consumption after DCZ relative to vehicle in *Gfral::hM4Di* animals. **i,** Representative images of stomach (white outline) and small intestine (white arrows) at selected time points following water barium contrast infusion, in *Gfral::ChR2* mice without (laser OFF) and with (laser ON) optogenetic stimulation. Laser was turned on 1 min before the start of the infusion. Scale bar: 10 mm. **j,** Left, fraction of maximum stomach area over time after infusion for laser off vs on. Right, maximum stomach area for each animal in laser off vs on. ns, not significant, *P<0.05, **P<0.01, ***P<0.001, ****P<0.0001. Data are mean ± SEM.

The activation of GFRAL neurons by fat ingestion was similar in magnitude to the response to high-dose emetics (e.g. 3.40 ± 0.65z for LiCl vs 3.54 ± 0.74z for Intralipid), yet animals showed no visible signs of distress, suggesting that GFRAL neuron activity might not be aversive in this context. We hypothesized that if Intralipid ingestion was aversive, then consumption would decline across repeated sessions as an aversive association is formed. Instead, Intralipid intake remained stable across four sessions (Fig. 3e), arguing against this interpretation. Of note, this is consistent with the extensive literature showing that mice find Intralipid appetitive^45–48^ and that it can condition flavor preferences^49–52^.

### GFRAL neurons are required for fat satiation and promote gastric accommodation

GFRAL neurons are known to suppress food intake^53–55^, which we confirmed by optogenetic stimulation of *Gfral::ChR2* mice (Extended Data Fig 4a-l). We therefore wondered whether the activation of GFRAL neurons by Intralipid consumption reflects a role in fat satiation.

To test this, we set out to silence GFRAL neurons and measure the effect on food intake. We generated *Gfral::hM4Di* mice and validated effective silencing in these animals by showing that deschloroclozapine (DCZ) treatment blocked conditioned taste avoidance induced by GDF-15 (0.36 ± 0.05 preference ratio for saline vs. 0.79 ± 0.05 preference ratio for DCZ, p=0.002; Fig. 3f). GFRAL silencing also partially rescued the anorexia induced by injection of LiCl (Extended Data Fig 4m-n). We then measured Intralipid and glucose consumption following vehicle or DCZ treatment. Silencing of GFRAL neurons significantly increased Intralipid consumption in hM4Di animals (447 ± 179 additional licks during DCZ vs. saline; Fig. 3g) but had no effect in control littermates (−392 ± 353 additional licks during DCZ vs. saline; Fig. 3g). Moreover, this effect was specific to fat, as glucose consumption was unaffected by GFRAL silencing in either group (Fig. 3h). Thus, endogenous GFRAL neuron activity is required for fat satiation at high levels of consumption.

In addition to promoting satiety, fat consumption also triggers gastric accommodation and inhibits gastric emptying, and GFRAL neuron activation is known to modulate gastric motility^53,54^. To visualize how GFRAL neuron activity modulates these GI reflexes, we used X-ray fluoroscopy^56^ to image gut dynamics in awake *Gfral::ChR2* mice (Extended Data Fig 5a). Animals received an intragastric infusion of a barium contrast reagent, with or without simultaneous optogenetic stimulation, and we quantified stomach contrast over time as a proxy for gastric emptying. GFRAL neuron stimulation caused a dramatic inhibition of gastric emptying, with most contrast reagent retained in the stomach 30 minutes after infusion (10.1 ± 6.8% retained at 30 min for light off vs. 85.5 ± 3.3% for light on, p=0.0043; Fig. 3i-j) as well as a significantly larger maximum stomach area compared to controls (71.1 ± 9.0 mm^2^ for laser-off vs. 208.2 ± 17.6 mm^2^ for laser-on, p= 0.0004; Fig. 3i-j).

This increase in stomach size following GFRAL neuron stimulation could reflect impaired gastric emptying, fundal relaxation, or a combination of both effects. To distinguish between these possibilities, we performed a second experiment where we allowed stomach filling to complete before turning on the laser. Remarkably, stomach size increased significantly within minutes of GFRAL neuron stimulation even in the absence of any additional infusion (93.6 ± 6.2% for laser-off vs. 135.4 ± 11.3% for laser-on at 5 min, p= 0.017; Extended Data Fig 5b-c), indicating that GFRAL neuron activation promotes active gastric expansion. Together, these findings establish that GFRAL neuron activity orchestrates a coordinated physiologic response to fat consumption that includes suppression of further intake, slowing of gastric emptying, and increase in gastric volume.

### Activation of GFRAL neurons by fat does not require canonical gut-brain pathways

We next investigated the mechanism by which GFRAL neurons are selectively activated by fat by blocking candidate gut-brain pathways. Fatty acids are sensed in the small intestine through the receptors GPR40 and GPR120 and the transporter CD36^57–59^, and combined blockade of these proteins prevents activation of many downstream brain circuits^47,51,60^. However, we found that a cocktail of antagonists against these proteins had no effect on GFRAL neuron activation by fat (2.02 ± 0.35z for vehicle vs. 1.48 ± 0.21z for inhibitors, p=0.39; Fig. 4a). Similarly, the peptides CCK and GLP1 are released from the intestine preferentially in response to fat, and are required for fat-induced activation of some hypothalamic and brainstem neurons involved in feeding^61,62^, but blockade of their receptors did not prevent GFRAL neuron activation by fat (2.04 ± 0.34 for vehicle vs. 2.39 ± 0.46 for devazepide, p=0.61, Fig. 4b; 2.43 ± 0.64 for vehicle vs. 2.08 ± 0.40 for ex-9-39, p=0.79 Fig. 4c). Finally, GDF-15 levels are known to rise in response to ingestion of some fats, albeit on a longer timescale than our recordings^63–65^. However, pretreatment with ponsegromab to neutralize GDF-15 did not alter GFRAL neuron responses to fat (2.89 ± 0.50 for vehicle vs. 2.67 ± 0.45, p=0.93; Fig. 4d). Thus, canonical gut-brain pathways for fat sensing appear to be dispensable for GFRAL neuron activation.

**Figure 4.**
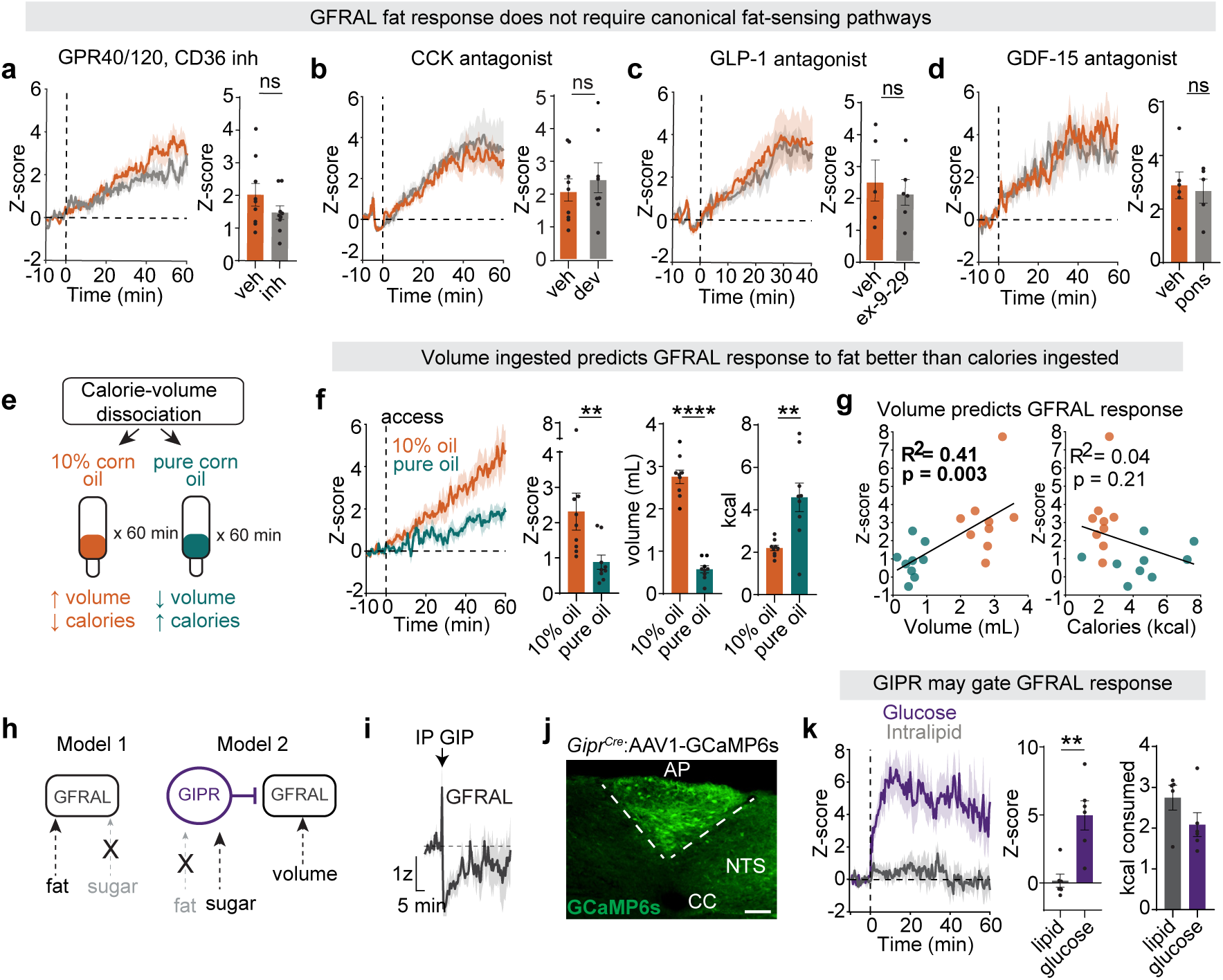
GFRAL neuron response to fat is related to ingested volume, and may be gated by inhibitory GIPR neurons. **a,** Left, Responses of GFRAL neurons to lipid ingestion with inhibitors (GW1100, AH7614, Sulfo-N-succinimidyl Oleate sodium) or vehicle. Right, Z-score bar plots. **b,** Animals received an IP injection of devazepide (CCK antagonist) or vehicle, 5 min prior to intralipid access. Left, Responses of GFRAL neurons to lipid after IP devazepide or vehicle. Right Z-score bar plots. **c,** Left, Responses of GFRAL neurons to lipid ingestion after IP exendin (9-39) or vehicle. Right, Z-score bar plots. **d,** Responses of GFRAL neurons to lipid ingestion after subq ponsegromab (GDF-15 neutralizing antibody) or vehicle. Right, Z-score bar plots. **e,** schematic of experimental design. Animals were allowed to lick either pure corn oil or 10% corn oil emulsion. **f,** Left, GFRAL neuron response to free licking of either a 10% corn oil emulsion or pure corn oil. Right, Z-score, volume consumed, and kcal consumed (in 60 min). **g,** Regression of z-score response as a function of volume or calories consumed. **h,** Schematic for possible models for GFRAL neuron fat selectivity. Left, GFRAL neurons have dedicated fat sensing pathways, but not glucose sensing pathways. Right, GFRAL neurons detect the presence of ingested volume, but a second inhibitory neuron (GIPR neurons in AP) responds specifically to sugar, and silences GFRAL neurons only during sugar but not fat consumption. **i,** Response of GFRAL neurons to IP injection of [D-Ala2]-GIP. **j,** Expression of GCaMP6s in GIPR neurons in the AP. scale bar: 100um. **k,** Left, GIPR neurons responses to lipid and glucose consumption. Right, z-score and volume ingested. ns, not significant. *P < 0.05, **P < 0.01, ****P < 0.0001. Data are mean +/- SEM

Given that GFRAL neuron responses scaled with cumulative fat consumed (Fig. 3d), we next tested the role of calories and volume. Mice were allowed to drink either pure corn oil or a 10% corn oil emulsion and GFRAL neuron responses were recorded (Fig. 4e). We found that, compared to pure corn oil, mice consumed a larger volume of the 10% emulsion (2.75 ± 0.16mL for emulsion vs. 0.57 ± 0.08mL for pure, p<0.0001; Fig. 4f) but fewer calories (2.20 ± 0.13kcal for emulsion vs. 4.59 ± 0.67kcal for pure, p=0.0036; Fig. 4f) due to the difference in caloric density. However, GFRAL responses were significantly larger during emulsion consumption (2.31 ± 0.53z for emulsion vs. 0.88 ± 0.20z for pure corn oil, p=0.0028, Fig. 4f). Regression analysis confirmed that ingested volume was a strong predictor of GFRAL neuron activation (R^2^ = 0.41, p = 0.003; Figure 4g), whereas calories consumed was not (R^2^ = 0.04, p = 0.21; Fig. 4g). Together, these data show that ingested volume is an important determinant of the GFRAL neuron response to fat.

### GIPR “anti-nausea” neurons are activated by sugar but not fat

While GFRAL neurons are activated in a volume-dependent manner when mice consume pure fat (Fig. 4f), we observed much less GFRAL activation when mice consumed similar volumes of a complete diet (Ensure: 30% fat; Fig. 3b-c). This raises the question of how GFRAL neuron activity can be both macronutrient and volume dependent. Given that GFRAL neurons were insensitive to blockade of common gut-brain pathways for fat (Fig. 4a-d), we considered the possibility that their responses to ingestion are instead shaped by an upstream neuron through feedforward inhibition (Fig. 4h, model 2). In this regard, GIPR-expressing neurons in the AP are GABAergic and have been shown to directly inhibit GFRAL neurons *ex vivo* and suppress GFRAL-activation induced aversion^7^. Therefore, if GIPR neurons are activated by sugar but not fat, this would give GFRAL neurons the appearance of fat specificity, even if they respond to a more general ingestive signal such as volume (Fig. 4h). This question is important, in part, because it remains mysterious how and when these “anti-nausea” GIPR neurons are naturally activated^66–68^.

To test this model, we first confirmed that activation of GIPR signaling does inhibit GFRAL neurons *in vivo*. GFRAL photometry mice were injected with the GIPR agonist [D-Ala2]-GIP, which produced a rapid suppression of baseline GFRAL activity that persisted for 34.0 ± 3.4 min (Fig. 4i), consistent with an inhibitory GIPR to GFRAL connection within the AP. We next generated mice expressing GCaMP6s in the AP of GIPR-Cre mice and recorded their activity during Intralipid and glucose consumption (Fig. 4j). Remarkably, GIPR neurons were robustly activated during glucose consumption but showed no response to Intralipid (4.97 ± 1.07z for glucose vs. 0.28 ± 0.57z for Intralipid, p=0.009; Fig. 4k), despite similar levels of consumption (2.09 ± 0.29kcal for glucose vs. 2.74 ± 0.30kcal for Intralipid, p=0.33; Fig. 4k). This reveals that these “anti-nausea” neurons^8,66–68^, which are key targets of the drug tirzepatide^69–71^, are activated during normal food ingestion, and suggests a model in which GIPR neurons gate GFRAL neuron activity in a macronutrient-specific manner (Fig. 4h).

### CALCR neurons are activated by food ingestion

The preceding data reveal that GFRAL and GIPR neurons are turned to the ingestion of fat and sugar, respectively. This surprising selectivity raises the question of whether other AP cell types track specific aspects of food ingestion. CALCR neurons, which are the target of an emerging class of weight loss medications exemplified by cagrilintide^72^, were the only cell type in our screen that responded robustly to Ensure consumption (Fig. 1). We therefore further investigated their natural activity during behavior.

We first characterized the response of CALCR neurons to self-paced consumption of Ensure (Fig. 5a-d). CALCR neurons showed ramping activation that began within a few seconds of the first lick, progressively increased several minutes (tau: 3.1 ± 0.8 min), and then remained elevated for the duration of the 40-minute recording, including during periods when animals were not licking (Fig. 5a-c, Extended Data Fig 6a,b). Consistent with this, decomposing CALCR neuron activity into fast (aligned to licks) and slow components revealed that the slow component contributed significantly more to the overall response than the fast component (3.73 ± 0.45z for slow vs. 1.79 ± 0.27 for fast, p=0.0078; Fig. 5d). This suggests that CALCR neurons respond to a post-ingestive signal that builds up during the first few minutes of ingestion and therefore likely arises from either the stomach or duodenum.

**Figure 5.**
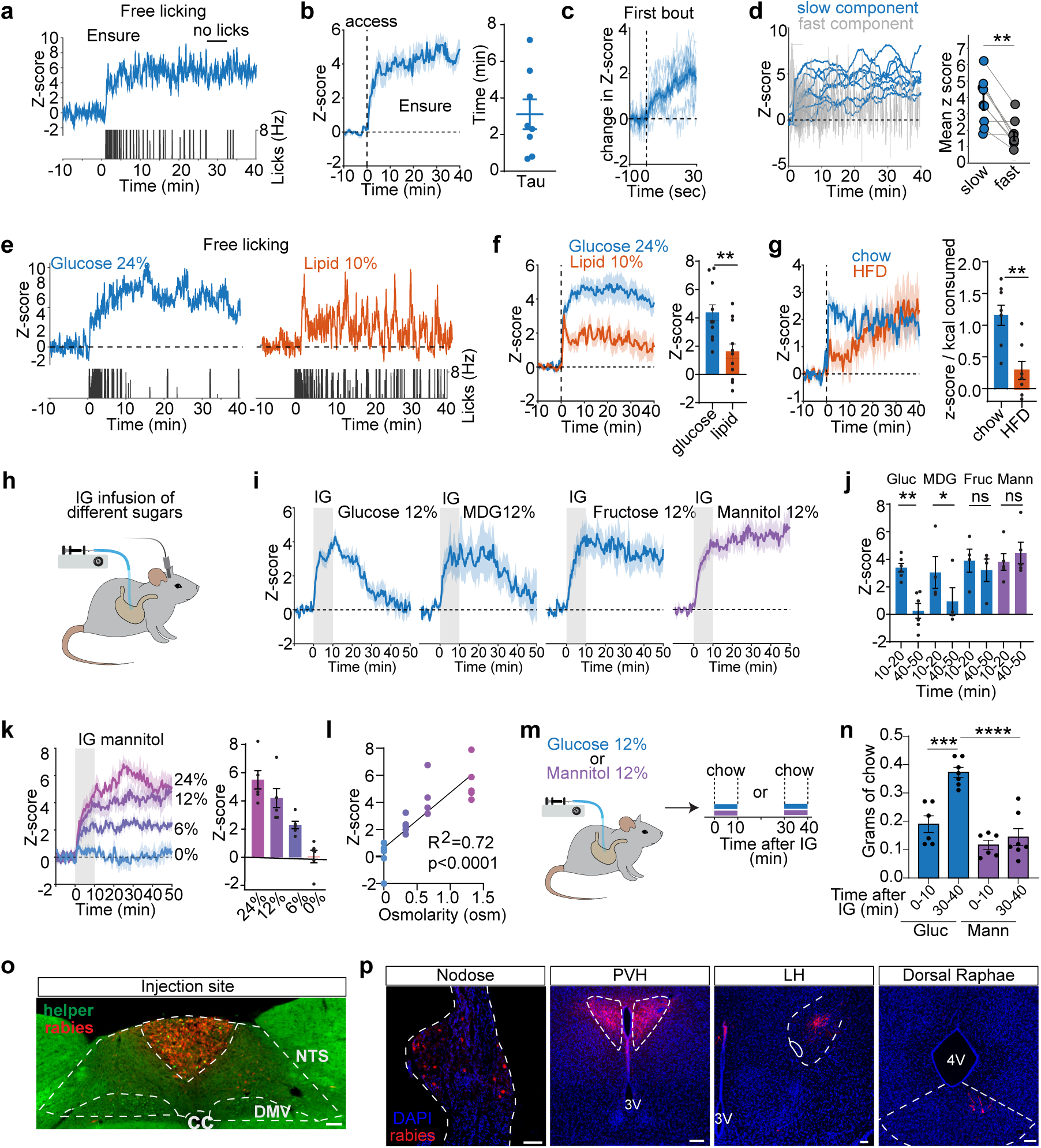
CALCR neurons encode ingestion through intestinal osmolarity. **a,** Example trace of calcium dynamics of CALCR neurons during self-paced Ensure consumption. The lick rate is shown below. **b,** Left, CALCR neuron activity aligned to access (dashed line) to liquid diet Ensure. Right, time constant (tau), when 63.8% of the z-scored activity change is reached. **c,** CALCR neuron responses aligned to the first lick of the first bout. **d,** CALCR activity was decomposed into fast (time locked to licking) and slow (changes in baseline z-score) components by fitting the signal to a saturating exponential and subtracting the slow fit from the raw trace. Left, fast and slow components for individual animals. Right, mean z score during slow vs fast component. **e,** Example traces of calcium dynamics during consumption of glucose vs lipid. **f,** Left, CALCR neuron responses aligned to self paced ingestion of glucose vs lipid. Right, z-score (0-40 min). **g,** Left, CALCR neuron responses aligned to access to chow vs high fat diet (HFD). Right, z-score normalized to caloric intake (0-40 min). **h,** schematic showing intragatric (IG) infusion of different sugars and sugar analogues. **i,** CALCR neuron responses to an intragastric infusion (0-10 min, 1.0 mL) of different sugar solutions. Gray bar shows IG infusion window. **j,** z-score for IG infusions in i, for early (10-20 min) vs late (40-50 min) time points after the start of the infusion. **k,** CALCR neuron responses to an intragastric infusion (0-10 min, 1.0 mL) of mannitol solutions of different concentrations. Right, z-score (0-40 min after infusion). **l,** Linear regression of CALCR z-score (mean 0-40 min after infusion) vs osmolarity of mannitol infused for data shown in k. Each dot is an individual animal colored by mannitol concentration. **m,** Experimental schematic, animals were given an IG infusion of either glucose 12% or mannitol 12%, and then received access to chow either 0-10 min after the infusion or 30-40 min after the infusion. **n,** Grams of chow consumed for each infusion type and time window. **o,** Expression of helper viruses (green) is restricted to the AP. Rabies virus expression in AP (red). **p,** Monosynaptic inputs onto CALCR neurons include nodose ganglion, paraventricular hypothalamus (PVH), lateral hypothalamus (LH), and dorsal raphae. scale bar: 100um. ns, not significant,*P<0.05, **P<0.01, ***P<0.001 ****P<0.0001. Data are mean ± SEM.

To determine which properties of food drive this response, we compared the response to consumption or intragastric (IG) infusion of different macronutrients. We found that CALCR neurons were activated during consumption of both glucose and Intralipid, albeit with stronger responses to sugar (4.37 ± 0.55z during glucose consumption vs 1.63 ± 0.51z during lipid consumption, p=0.0037; Fig. 5e,f). We observed similar results with calorie and volume-matched IG infusions of the same solutions (3.53 ± 0.63z during glucose infusion vs. 0.99 ± 0.77z during lipid infusion, p=0.028; Extended Data Fig 6c). Consistently, CALCR neurons were activated during consumption of both chow and high fat diet (HFD), but the response to chow was faster and larger (1.16 ± 0.15z/kcal for chow vs. 0.29 ± 0.15z/kcal, p=0.004; Fig. 5g) despite consumption of nearly three times more HFD. These results indicate that CALCR neurons are broadly activated during ingestion, with a bias toward foods high in carbohydrates.

### CALCR neurons track intestinal osmolarity

The kinetics of CALCR neuron activation during ingestion are too slow to be primarily orosensory, and, consistently, we found that consumption of the non-caloric sweetener sucralose caused minimal activation (3.30 ± 0.24z for glucose vs. 1.01 ± 0.24z for sucralose, p=0.0004; Extended Data Fig 6d). Moreover, this activation, unlike for glucose, was not sustained after access was removed (2.05 ± 0.45z for glucose vs. 0.43 ± 0.52z, p=0.0046; Extended Data Fig 6d). This indicates that sustained CALCR neuron activation requires a post-ingestive signal.

To identify what is sensed in the gut to drive CALCR neuron activation, we compared the response to IG infusion of volume and concentration-matched solutions of four sugars or sugar analogs that differ in their caloric content, absorption kinetics, and detection mechanism: glucose, fructose, a non-caloric SGLT1 agonist (MDG), and a non-caloric sugar alcohol (mannitol) (Fig. 5h-j). All four solutions activated CALCR neurons robustly during the 10 min infusion. However, these solutions differed greatly in how long CALCR neuron activation persisted after the infusion ended (Fig. 5i,j), and this differential decay mirrored their respective absorption kinetics. That is, glucose and MDG, which are rapidly cleared from the intestinal lumen^73,74^, elicited CALCR neuron responses that declined rapidly after the infusion ended, whereas responses to fructose (which is absorbed slowly^73^) and mannitol (which is not absorbed at all^74^) were much more sustained (Fig. 5i,j). Together, these results indicate that neither calories nor SGLT1 activation is required for CALCR neuron activation. Rather, these neurons appear to track the osmolarity of the luminal contents of the intestine, which remains elevated for as long as the sugar is present in the intestine.

To directly test the role of luminal osmolarity, we infused animals with mannitol solutions of varying concentrations (0%, 6%, 12%, and 24%) and measured CALCR neuron responses (Fig. 5k). This revealed that the response magnitude was linearly correlated with the osmolarity of the infused solution (R^2^ = 0.72, p < 0.0001; Fig. 5l). Similarly, higher concentrations of glucose, MDG, and fructose also produced progressively larger CALCR neuron responses (Extended Data Fig 6e-g). This indicates that CALCR neurons respond to intestinal osmolarity, and that the preferential activation by sugar over fat likely arises because sugar solutions are substantially more hyperosmotic than isocaloric fat emulsions.

Hyperosmolar fluids in the intestine are known to induce intestinal stretch, which activates vagal sensory neurons and thereby inhibits food intake^75–77^. We therefore hypothesized that the time course of CALCR neuron activation by intestinal hyperosmolarity would correlate with the time course of the suppression of feeding. To test this, we took advantage of the fact that glucose activates CALCR neurons early after infusion but decays as the sugar is absorbed, whereas mannitol induces elevated activation at both early and late time points (Fig. 5i,j). Animals were food deprived overnight, received an IG infusion of either glucose 12% or mannitol 12%, and were then given access to chow either 0-10 min or 30-40 min after infusion (Fig. 5m). At the early time point, both glucose and mannitol similarly suppressed chow intake, whereas at the later time point, animals that received glucose ate significantly more than animals that received mannitol (0.37 ± 0.02g for glucose vs. 0.14 ± 0.03g for mannitol, p<0.0001; Fig. 5n), consistent with the differential dynamics of CALCR neuron activation by these two substances. These data indicate that CALCR neuron dynamics correlate with feeding suppression.

The fact that CALCR neurons are activated by intestinal hyperosmolarity suggests that they may get input from the vagus nerve^77^. To identify the neural inputs to CALCR neurons, we performed monosynaptic retrograde rabies tracing (Fig. 5o), which revealed labeled presynaptic cells in the nodose ganglion as well as neurons in the paraventricular hypothalamus (PVH), lateral hypothalamus (LH), and dorsal raphae, regions associated with the control of food intake (Fig. 5p). Thus, CALCR neurons are positioned to integrate ascending visceral signals from the vagus nerve with descending input from forebrain circuits controlling energy balance and feeding.

### CALCR neurons suppress feeding and delay gastric emptying

Given that CALCR neuron activation correlates with a decrease in feeding (Fig. 5n), we next tested whether stimulation of these cells is sufficient to suppress food intake. We targeted hM3Dq to CALCR neurons by injecting a Cre-dependent AAV into the AP and confirmed that this did not infect the adjacent NTS (Fig. 6a). Activating CALCR neurons decreased consumption of Ensure (2941 ± 224 licks for vehicle vs. 1646 ± 257 licks for DCZ, p=0.0020; Fig. 6b) through a reduction in the number of bouts, with no change in bout duration (37 ± 2 bouts for vehicle vs. 23 ± 3 bouts for DCZ, p=0.009; Fig. 6c). Activating CALCR neurons also strongly inhibited chow consumption in fasted animals and water consumption in thirsty animals (Fig. 6d, Extended Data Fig 7a,b). Notably, the degree of feeding inhibition correlated with the level of hM3Dq expression in the AP quantified by post-hoc histology (Extended Data Fig 7c-e). Moreover, CALCR neuron stimulation induced FOS expression in downstream brain regions that promote satiety (Fig. 6e,f), consistent with a role for CALCR neuron activity in inhibiting ingestion^24,78^.

**Figure 6.**
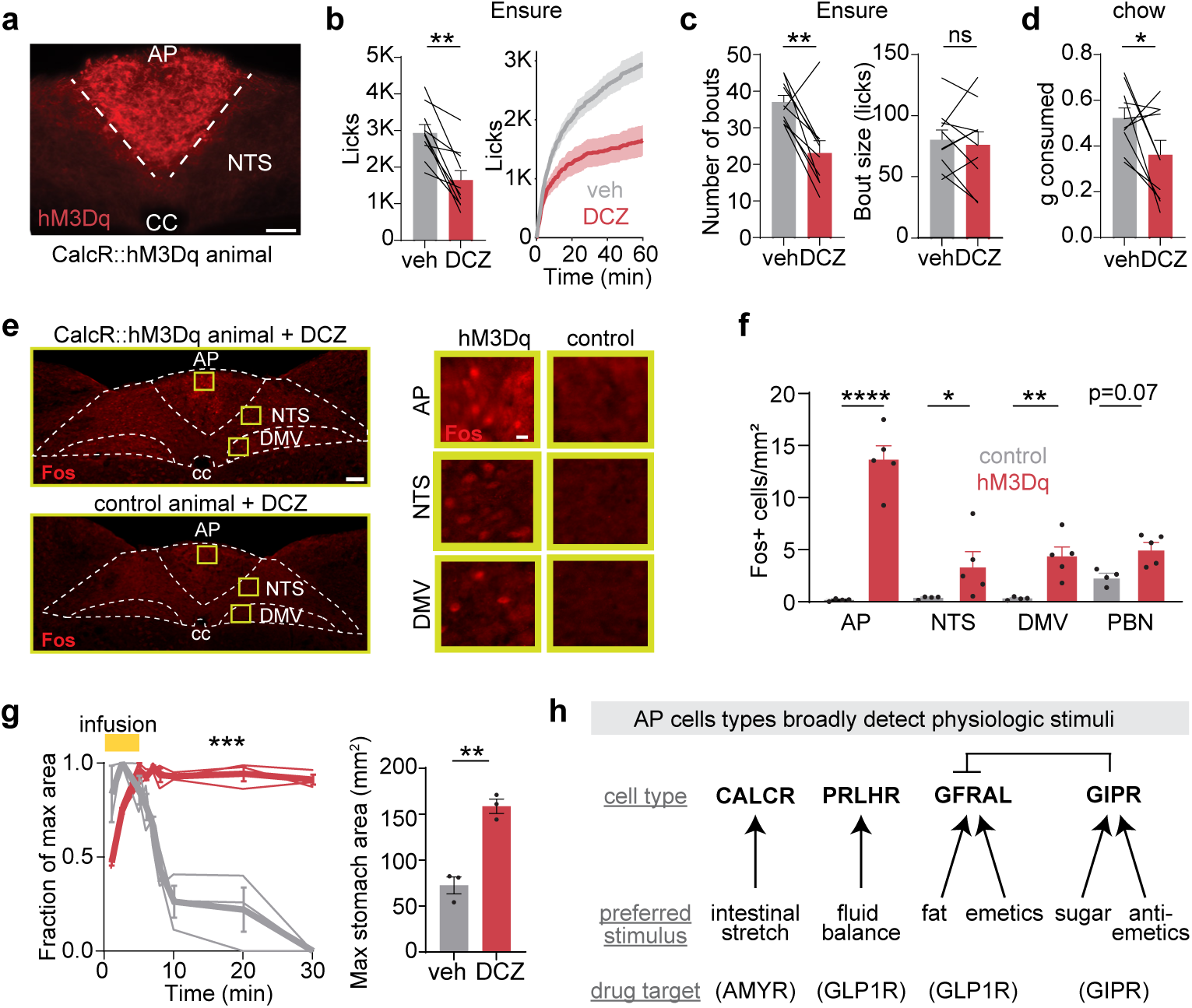
CALCR neurons pace food ingestion and slow down gastric emptying. **a,** Expression of hM3Dq::mCherry restricted to the area postrema (AP) without expression in adjacent nucleus tractus solitarius (NTS). cc: central canal, scale bar: 100um. **b,** Ensure consumption (during 60 min) after either vehicle or DCZ injection in animals expressing hM3Dq::mCherry in CALCR AP neurons. Left, total number of licks. Right, cumulative licks. **c,** Left, number of bouts. Right, bout size (licks). **d,** Chow consumption after either veh or DCZ injection. **e,** Fos expression in dorsal vagal complex in either experimental hM3Dq::mCherry or control animals, 2 hrs after DCZ injection. DMV:dorsal motor vagus. **f,** Fos expression levels in select brain regions. **g,** Left, Fluoroscopy experiment showing fraction of stomach maximum area over time since the infusion. Right, maximum stomach area at any time during the infusion. **h,** Schematic summarizing the findings of this study. Genetically-defined AP cell types are tuned to different aspects of physiology. **\***P<0.05, **P<0.01, ***P<0.001 ****P<0.0001. Data are mean ± SEM.

CALCR neurons stimulation also induced FOS in the dorsal motor vagus (Fig. 6e,f), which contains the cell bodies of vagal efferents that control GI motility. To examine how CALCR neurons control GI dynamics, we stimulated CALCR neurons in awake *Calcr::hM3Dq* mice and then imaged the gut by X-ray fluoroscopy. In vehicle-treated animals, barium contrast cleared rapidly from the stomach, with near-complete emptying within 30 minutes. In contrast, DCZ-treated animals showed a dramatic slowing of gastric emptying, with barium contrast largely retained throughout the imaging session (0.0 ± 0.0% retained at 30 min for vehicle vs. 91.2 ± 2.7% for DCZ, p=0.0002; Fig. 6g) and a significantly larger maximum stomach area compared to controls (72.5 ± 9.2mm^2^ for vehicle vs. 158.8 ± 7.9mm^2^ for DCZ, p=0.0023; Fig. 6g, Extended Data Fig 7f). Altogether, these findings reveal that CALCR neurons are activated by hyperosmotic solutions entering the intestinal lumen and, in turn, coordinate two counterregulatory responses: suppression of ongoing food intake and slowing of gastric emptying, both which reduce the rate at which further osmotic loads are delivered to the intestine.

## Discussion

The area postrema was identified 75 years ago as the “chemoreceptor trigger zone” responsible for nausea in response to circulating toxins and emetics^1,2^. Nevertheless, the recent discovery that AP neurons are the direct targets of GLP-1 drugs, and appear to mediate much of their weight loss effects^9–12^, has stimulated interest in the AP’s broader role in non-aversive physiology. Here, we have investigated this question by recording the activity of an array of AP cell types during behavior, including the cells that are the principal targets of each major class of peptide hormone drugs used for obesity. We find that each of these cell types exhibits strong and specific activation in response to a different physiologic stimulus that is unrelated to sickness (Fig. 6h). We show that GFRAL neurons, the brain’s prototypical nausea neuron, are selectively activated by consumption of fat and are required for fat satiation, whereas GIPR-expressing “anti-nausea” neurons are selectively activated by sugar. CALCR neurons sense intestinal osmolarity during food ingestion and trigger a coordinated response that slows the delivery of osmotic loads to the intestine, whereas PRLHR neurons are activated by hypovolemia via an angiotensin-dependent mechanism, suggesting that they are responsible for the AP’s role in blood pressure regulation^79–81^. Together, these findings reveal that genetically-defined AP cell types delineate parallel pathways for sensing and responding to a broad range of physiologic stimuli unrelated to sickness and aversion (Fig. 6h).

### Nausea and anti-nausea neurons are tuned to ingestion of fat and sugar, respectively

GFRAL neurons are activated by GDF-15^25–27^ and, for this reason, have been thought to have a dedicated function in nausea and sickness. In this context, it was surprising to discover that GFRAL neurons are strongly activated by consumption of fat, which is not normally aversive. Nevertheless, there is some previous evidence suggesting GFRAL neurons play a role in fat metabolism. This includes the fact that (1) consumption of fats leads to increased GDF-15 levels in the blood^63–65^, albeit on longer timescales than the response we observe by photometry, (2) GDF-15 and GFRAL knockout mice have exacerbated obesity when placed on a high-fat diet^26,82,83^, and (3) stimulation of GFRAL neurons promotes a shift from glucose to fatty acid oxidation in peripheral tissues^25,54,55,84^. These findings have been interpreted in the context of sickness and chronic inflammation, but our data suggest that GFRAL neurons may have a more basic role in sensing and responding to lipid ingestion.

Despite strong activation of GFRAL neurons during fat consumption, we saw no evidence that this was aversive, consistent with the extensive literature showing that consumption of fats such as Intralipid and corn oil is rewarding^45–48^ and conditions flavor preferences^49–51^. We do not know the basis for this apparent context-dependent valence of GFRAL neuron activity. One possibility is that the valence of GFRAL activation depends on aspects of neural dynamics not apparent from population recordings by photometry. An alternative possibility is that food consumption generates signals that act downstream of GFRAL neurons (e.g. in the parabrachial nucleus) to suppress their effects on nausea but not satiety.

GFRAL neurons are directly inhibited by neurons expressing GIPR, and this circuit is thought to be responsible for the anti-nausea effects of GIPR agonists including tirzepatide^8,66–68^. It has been unknown, however, under which natural circumstances these “anti-nausea” neurons become activated. We found surprisingly that GIPR neurons are strongly activated by consumption of sugar but not pure fat. This is unlikely to be mediated by the peptide GIP, which is released at higher levels in response to fat compared to sugar^85,86^. The activation of GIPR neurons by sugar also suggests that consumption of sugar solutions alone should have an anti-emetic effect. While counterintuitive, this is consistent with the fact that a highly concentrated sugar solution, known as Emetrol, has long been marketed for nausea and has been shown to have anti-nausea effects in clinical trials^87–89^. It is possible that this over-the-counter anti-emetic recruits the same GIPR brainstem circuit that is targeted by tirzepatide.

### CALCR neurons detect intestinal stretch to promote non-aversive satiety

The CALCR is the target of an emerging class of anti-obesity therapeutics that include cagrilintide^78,90^. While it is known that CALCR is the receptor for amylin^91^, a hormone released from the pancreas along with insulin, it has been unknown when and how CALCR neurons are naturally activated.

Here, we found that CALCR cells are activated by the increase in luminal osmolarity that occurs when nutrients enter the small intestine, and their activation in turn suppresses food intake and slows down gastric emptying. This activation was directly proportional to osmolarity and did not depend on caloric value of the substance ingested, suggesting it is unlikely to be mediated by nutrient-sensitive hormones, including amylin. Rather, the response likely involves intestinal stretch, which is induced by influx of water into the intestine following a hyperosmotic load and is detected by OXTR-expressing vagal afferents, which like CALCR neurons, potently inhibit food intake when stimulated^77,92^. In a natural setting, this stretch-sensitive pathway might be engaged most strongly when animals consume foods high in fructose, such as fruit, since this sugar has a high osmotic density per calorie and is slowly cleared from the intestine.

### The area postrema can detect physiologic stimuli through rapid mechanisms

The AP is a circumventricular organ and for this reason is traditionally assumed to detect slow changes in circulating hormones and nutrients^4–6^. However, we find that AP dynamics during behavior are dominated by rapid responses that are unlikely to be mediated by circulating signals. For example, we found that emetics rapidly activate GFRAL neurons by a mechanism independent of GDF15. This was observed even for cisplatin, which induces robust GDF15 expression^32,39^. Similarly, GFRAL neuron activation by fat ingestion did not require GDF15 or known fat-sensitive hormones, and CALCR neuron activation during ingestion was calorie-independent and therefore likely due to intestinal stretch. Consistently, we found that both cell types receive abundant monosynaptic input from the forebrain and vagus nerve, providing a pathway for rapid modulation of their activity. It is likely that the natural role of many hormones is to modulate AP activity on a longer timescale than our photometry recordings. An important challenge for the future will be to understand how these fast and slow signals are integrated in the AP to control physiology and behavior.

## Acknowledgements

We thank members of the Knight laboratory for discussions. We thank J. Grove, B. Jarvie, and J. Jimenez for comments on the manuscript. We thank M. Andermann for advice on x-ray fluoroscopy. This work was supported by NIH grants R01-DK106399, R01-DK138127, and R01-DK145100 (Z.A.K.), and T32DK007007 and HHMI GT17734 (A.L.C). A.L.C is an HHMI Hanna Gray Fellow. Z.A.K. is an Investigator of the Howard Hughes Medical Institute.

## Author Contributions

A.L.C and Z.A.K. conceived the project and designed experiments. A.L.C. led the experiments and analyzed the data. A.L.C, K.X., M.M, M.K, A.S., and K.C. performed photometry and chemogenetic experiments. A.L.C, and K.X. performed optogenetic experiments. N.F.B and A.L.C conducted rabies tracing experiments. A.M.H. and A.L.C conducted Fos experiments. A.L.C. conducted all fluoroscopy experiments. A.L.C performed all surgeries except intragastric catheters. N.F.B and A.E.A. provided guidance and training on brainstem injections. L.Q. and A.M.H. performed intragastric catheter surgeries. A.L.C, N.F.B, A.M.H., M.M, and M.K. collected and analyzed histology. T.E.L provided guidance and training on fluoroscopy experiments. F.M.G., F.R., and M.G.M generated transgenic mice. A.L.C and Z.A.K. wrote the manuscript with input from all authors.

## Additional information

Correspondence and requests for materials should be addressed to Zachary A. Knight.

## Data availability

The data from this study are available from the corresponding author on reasonable request.

## Methods

All experimental protocols were approved by the Institutional Animal Care and Use Committee of the University of California, San Francisco, following the National Institutes of Health guidelines for the Care and Use of Laboratory Animals.

### Mouse strains

Experimental animals (>8 weeks old, both sexes) were kept in temperature-controlled and humidity-controlled facilities with a 12-h light-dark cycle and with ad libitum access to water and standard chow (PicoLab, 5053). The following mice were obtained from the Jackson Laboratory: WT (C57BL/6J; 000664); Calcr-cre (STOCK Calcrtm1.1(cre)Mgmj/J, 037028), Gfral-cre (B6;SJL-Gfralem1(cre)Rsy/J, 036750), Slc6a2-cre (C57BL/6J-Slc6a2em1(cre)Lbrl/J, 037882), TIGRE2-jGCaMP8s-IRES-tTA2 (STOCK Igs7tm2(tetO-GCaMP8s,CAG-tTA2)Genie/J, 037719), Ai32 (B6.Cg-Gt(ROSA)26Sortm32(CAG-COP4*H134R/EYFP)Hze/J, 024109), and R26-LSL-Gi-DREADD (B6.129-Gt(ROSA)26Sortm1(CAG-CHRM4*,-mCitrine)Ute/J, 026219). Gipr-cre mice were a gift from Frank Reimann. Prlhr-cre mice were a gift from Martin Myers. Gfral-cre, Calcr-cre, Prlhr-cre, and Slc6a2-cre mice were crossed to jGCaMP8s-IRES-tTA2 mice to generate double mutants (Gfral-Cre/+GCaMP8s/+, Calcr-Cre/+GCaMP8s/+, Prlhr-Cre/+GCaMP8s/+, Slc6a2-Cre/+GCaMP8s/+ mice, respectively). Gfral-Cre mice were crossed to Ai32 and R26-LSL-Gi-DREADD mice to generate double mutants (Gfral-Cre/+RosaChR2/+ and Gfral-Cre/+hM4Di/+, respectively).

### Intracranial surgery

#### General procedures

##### For implants

Animals were anesthetized with 2% isoflurane and secured in a stereotaxic frame on a heating pad. Ophthalmic ointment was applied to protect the eyes, and subcutaneous injections of meloxicam (5.0 mg/kg) and sustained-release buprenorphine (1.5 mg/kg) were administered prior to surgery. The scalp was shaved and scrubbed with betadine and alcohol (three times), local anesthetic was applied (bupivacaine 0.25%), and a midline incision was made. A small craniotomy was performed using a dental drill (0.5 mm burr hole). Fiber optic cannulas were then implanted and secured to the skull with Metabond (Patterson Dental Supply, 07-5533559, 07-5533500; Henry Schein, 1864477) and Flow-It (Patterson Dental Supply, 07-6472542). To reduce postoperative inflammation, animals received dexamethasone (0.6 mg/kg) for three days following surgery.

##### For AP viral injections

Animals were anesthetized with ketamine (10 mg/kg) and xylazine (1 mg/kg). Ophthalmic ointment was applied to the eyes and a subcutaneous injection of meloxicam (5.0 mg/kg) was given prior to the surgery. The scalp was shaved, scrubbed (betadine and alcohol three times), and local anesthetic applied (bupivacaine 0.25%). Animals were placed in the stereotactic frame on a heating pad, and the head was tilted forward 90 degrees relative to the body. An incision was made below the occipital crest, the neck muscles were retracted, the meninges were dissected, and the caudal brainstem was exposed and directly visualized. Virus was injected at a rate of 50 nL/min using a beveled glass pipette connected to a 10 µl Hamilton syringe (WPI), controlled using a Micro4 microsyringe pump controller (WPI). The needle was kept at the injection site for 1 min before withdrawal. After injection, retractors were removed, the skin was sutured and animals were given dexamethasone (0.6 mg/kg) and sustained-release buprenorphine (1.5 mg /kg)

### Fiber photometry implants in the AP

Gfral-Cre/+GCaMP8s/+, Calcr-Cre/+GCaMP8s/+, Prlhr-Cre/+GCaMP8s/+, Slc6a2-Cre/+GCaMP8s/+, and Gipr-Cre/AAV1GCaMP6s mice were prepared for photometry recordings by implanting an optic fiber (Doric Lenses, MFC_400/430-0.48_6mm_MF2.5_FLT) and sleeve (Doric Lenses, SLEEVE_BR_2.5) above the area postrema (1.5 mm anterior–posterior, 0 mm medial–lateral (ML) and −4.05 mm dorsal–ventral (DV), relative to the occipital crest with 20° in the anterior-posterior direction). Mice were allowed to recover for a minimum of 3 weeks before the first photometry experiment. In separate surgeries, mice were equipped with an intragastric catheter.

### Viral injection into the AP

#### DREADDS virus injection

Calcr-Cre animals were prepared for viral injections as described above. The caudal brainstem was exposed and AAV-hSyn-DIO-hM3D(Gq)-mCherry (50 nl; 5.0 × 10^12^ viral genome copies (vg) per ml; Addgene 44361-AAV9) was injected into the area postrema (0.65 mm anterior–posterior (AP), 0 mm medial–lateral (ML) and −0.16 mm dorsal–ventral (DV) relative to the obex). Mice were allowed to recover for a minimum of 3 weeks before the first experiment. For fluoroscopy experiments head bars were affixed to the skull using Metabond and mice were equipped with an intragastric catheter in separate surgeries.

#### GCaMP6s virus injection

Gipr-Cre animals were prepared for viral injections as described above. The caudal brainstem was exposed and AAV1-CAG-Flex-GCaMP6s-WPRE-SV40 (50-100 nl; 3.0 × 10^12^ viral genome copies (vg) per ml; Addgene 100842-AAV1) was injected into the area postrema (0.65 mm anterior–posterior (AP), 0 mm medial–lateral (ML) and −0.16 mm dorsal–ventral (DV) relative to the obex). Mice were allowed to recover for a minimum of 1 week and then a photometry fiber was implanted in a second surgery.

#### Rabies tracing experiments

Calcr-Cre and Gfral-Cre animals were prepared for viral injections as described above. The caudal brainstem was exposed and the helper viruses AAV1-TREtight-mTagBFP2-N2cG (50 nl; 1.6 × 10^12^ viral genome copies (vg) per ml; Addgene192838) and AAV1-syn-FLEX-splitTVA-EGFP-tTA (50 nl; 8.5 × 10^10^ viral genome copies (vg) per ml; Addgene100798) were injected into the area postrema (0.65 mm anterior–posterior (AP), 0 mm medial–lateral (ML) and −0.16 mm dorsal–ventral (DV) relative to the obex). Mice were allowed 2 weeks for viral expression, and were subsequently prepped for a second surgery, in which RabV-CVS-N2c(deltaG)-mCherry(EnvA) (100 nl; 1.63 × 10^9^ viral genome copies (vg) per ml; Janelia) was injected into the area postrema (0.65 mm anterior–posterior (AP), 0 mm medial–lateral (ML) and −0.16 mm dorsal–ventral (DV) relative to the obex). Mice were allowed 7 days for viral expression/spread, prior to perfusion and tissue processing.

### Optogenetic implants in the AP

Gfral-cre/+, Gfral-cre/+RosaChR2/+, mice were prepared for optogenetic experiments by installing a fiber optic cannula (Doric, MFC_200/240-0.22_6mm_ZF1.25(G)_FLT) above the AP area postrema (1.5 mm anterior–posterior, 0 mm medial–lateral (ML) and −3.95 mm dorsal–ventral (DV) relative to the occipital crest with 20° in the anterior-posterior direction). Mice were allowed to recover for a minimum of 2 weeks before optogenetic experiments. For fluoroscopy experiments, head bars were affixed to the skull using Meta bond and intragastric catheters were placed in a separate surgery.

### Intragastric catheter surgery

Mice were deeply anesthetized with 2% isoflurane and surgical sites were shaved and sterilized with alternating betadine and ethanol scrubs. Meloxicam (5 mg/kg) and sustained-release buprenorphine (1.5 mg/kg) were administered subcutaneously prior to surgery. A midline abdominal incision of approximately 1.5 cm was made caudal to the xyphoid process, and a secondary 1 cm incision was made between the scapulae for catheter externalization. Blunt dissection was used to separate the skin from the subcutaneous tissue along the left flank, creating a tunnel between the two incisions for catheter routing. A small incision was made in the abdominal wall, and the catheter (Instech, C30PU-RGA1439) was guided through the interscapular incision into the abdominal cavity using curved hemostats. The stomach was gently externalized with atraumatic forceps and a purse-string suture was placed in the forestomach using 7-0 non-absorbable Ethilon. A small puncture was made at the center of the purse string, the catheter tip was inserted, and the suture was tightened to secure it in place, with 2–5 mm of catheter fixed within the stomach lumen. The abdominal cavity was irrigated with 1 mL of sterile saline and the stomach was returned to its anatomical position. The abdominal wall was closed in two layers, the catheter was anchored to the muscle at the interscapular site, and the interscapular incision was closed. The external catheter end was capped with a 22-gauge PinPort (Instech, PNP3F22). Mice received Baytril (5 mg kg⁻¹) and warm saline at the end of surgery and were allowed to recover for one week before the start of photometry experiments.

### Fiber photometry recordings

#### Photometry setup

Mice were connected to a patch cable (Doric Lenses, MFP_400/460/900-0.48_2m_FCM-MF2.5) for photometry recordings. A blue LED (470 nm) and a UV LED (405 nm) served as excitation sources for the calcium-dependent GCaMP signal and the isosbestic control signal, respectively. Both LEDs were driven by a multichannel hub (Thorlabs) and sinusoidally modulated at 305 Hz and 505 Hz before being delivered through a filtered minicube (Doric Lenses, FMC6_AE(400-410)_E1(450-490)_F1(500-540)_E2(550-580)_F2(600-680)_S) and through the implanted optic fiber. Emitted fluorescence was collected back through the same fiber, separated at the dichroic ports of the minicube, and detected by a femtowatt silicon photoreceiver (Newport, 2151). The resulting signals were digitally sampled at 1.0173 kHz, demodulated by lock-in amplification, and acquired through a real-time processor (RZ5P, Tucker-Davis Technologies) using Synapse software (TDT). Data was exported using TDT Browser and downsampled to 4 Hz in MATLAB prior to analysis.

#### Behavior

For all recordings, mice were placed in isolated behavioral chambers (Med Associate) without water or food access unless otherwise specified. Chambers were cleaned between experiments with 70% ethanol to remove olfactory cues from previous experiments. Mice were habituated for one night in the chambers before experiments with lickometers containing water and chow in the chamber. Before each recording, photometry implants on individual mice were cleaned with 70% ethanol using connector cleaning sticks (MCC-S25) and connected to a photometry patch cable immediately afterwards. For all photometry experiments, mice were acclimated to the behavioral chamber for at least 20 min with recording before presentation of a stimulus. For injection experiments, mice were injected with compounds at the following concentrations based on previously published reports: amylin rat, 20 μg/kg (MedChemExpress); salmon calcitonin, 150 μg/kg (MedChemExpress); ghrelin, 2 mg/kg (R&D Systems); [D-Ala2]-GIP, 374 μg/kg (Biotechne); LiCl, 168 mg/kg; Deoxynivalenol (DON), 2.5 mg/kg (Tocris); exendin-4, 100 μg/kg (Bachem), 3M NaCl; angiotensin-II, 2.0 mg/kg (MedChemExpress); cisplatin 7.5 mg/kg (MedChemExpress), GDF-15, 25 μg/kg (MedChemExpress); devazepide, 1 mg kg/1 (R&D Systems); Exendin(9-39), 1 mg/kg (MedChemExpress); ponsegromab, 10 mg/kg, (MedChemExpress), losartan, 100 mg/kg (MedChemExpress); captopril, 50 mg/kg (MedChemExpress); isoproterenol 100 mg/kg (MEdCHemExpress). All these compounds were dissolved in PBS, except devazepide, which was dissolved in 5% DMSO, 5% Tween-80 in saline (vehicle solution for devazepide). All compounds were injected at a volume of 10 µL/g of mouse body weight. For all ponsegromab experiments, mice received a subcutaneous injection 16 hours before the experiment.

For PRLHR experiments, losartan, captopril, and PEG (40%) were administered subcutaneously, and isoproterenol was administered intraperitoneally. To block angiotensin signaling, losartan was administered 20 min prior to angiotensin-II, and losartan + captopril (described “inh”) were administered 20 min prior to PEG. Losartan is a selective angiotensin type 1 receptor (AT1R) antagonist, and captopril is an angiotensin converting enzyme (ACE) inhibitor.

For the licking experiments, mice were given access to the tested solution in their home cage a few days before the experiment to avoid neophobia. Mice were fasted overnight and on the experimental day, mice received access to a lickometer containing the indicated solution for the time window of the experiment. Solutions were prepared using deionized water at the following concentrations: Ensure Original Vanilla Nutrition Shake with Fiber (Abbott Nutrition, undiluted), 0.12-0.24 g/mL glucose (12-24%), 0.12-0.24 g/mL fructose (12-24%), 0.12-0.24 g/mL methyl-D-glucopyranoside (12%-24% MDG), and 0.8 mg/mL sucralose (in PBS). Intralipid 20% (Sigma, I141-100ML; Medline, BHL2B6064H) was diluted 1:1 in water to make Intralipid 10%. Corn oil emulsion (10%) was made in deionized water with 0.1% xantham gum and 0.05% Tween 80 and blended for 5 min to emulsify. For chow and HFD experiments, mice were deprived of food overnight before the experiment. Mice were then given access to standard chow (PicoLab 5053) or a HFD (Research Diets, D12492) for the duration indicated in the experiment.

For i.g. infusion experiments, mice were deprived of food overnight before the experiment. Solutions, Ensure, glucose (12%, 24%), Intralipid (10%), MDG (12%, 24%), fructose (12, 24%) mannitol (6%, 12%, 24%), and water, were delivered using a syringe pump (Harvard Apparatus, 70–2001) over 10 min. The infusion rate was 0.1 mL/min for a total infusion volume of 1.0 mL. Before mice were placed into the Med Associates behavioral chambers for habituation, the i.g. catheter was attached to the syringe pump using plastic tubing and adapters (AAD04119, Tygon; LS20, Instech). Mice received an infusion of water and an infusion of Ensure for habituation, several days prior to the first experiment.

For Intralipid and inhibitor experiments: Animals were treated with an IP injection of devazepide or exendin(9-39) 5 min prior to access to the lickometer, and control animals received vehicle injection. Animals were treated with a subcutaneous injection of ponsegromab 16 hours before the Intralipid licking experiment, and control animals received vehicle injection. For blockade of GPR40,120, CD36, the following compounds were added to 10% Intralipid in DMSO: 50 µg/mL GW1100, 50 µg/mL AH761, and 10 µg/mL Sulfo-N-succinimidyl oleate (1.1% volume of inhibitors in DMSO). Control experiments were Intralipid 10% Intralipid plus 1.1% DMSO.

#### Analysis

GCaMP8s calcium signals acquired at 470 nm were normalized to the isosbestic control signal acquired at 405/415 nm using a linear regression fit derived from the baseline period, yielding a normalized fluorescence trace (Fnormalized). This normalized signal was then converted to a z-score using the mean and standard deviation of the baseline period preceding each stimulus: z = (Fnormalized − μ) / σ. For display purposes, mean traces were downsampled by a factor of 120 to reduce file size.

For most experiments, baseline activity was calculated from the 10 minutes preceding stimulus presentation, during which animals remained undisturbed in the behavior chamber. The mean z-score during this baseline window was compared to the average z-score during the post-stimulus epoch of interest. Mean z-scores reported are for the entire post stimulus epoch, unless otherwise noted.

To calculate the response time and T50 for each animal, z-scored photometry traces were smoothed using two lowess filters of different bandwidths: a heavy smooth (120 s window) for response time, and a moderate smooth (30 s window) for T50 crossing precision. Response time was defined as the first time point at which the heavily smoothed trace crossed zero and remained above zero for the remainder of the recording. T50 was defined as the first time point at which the moderately smoothed trace exceeded 50% of the peak z-score (determined from the heavily smoothed trace) and remained above this threshold for at least 10 consecutive seconds.

For GFRAL Intralipid responses, we performed single-variable linear regression between the z-scored photometry signal and three licking predictors: instantaneous lick rate, licks in the past 10 seconds, and cumulative licks from the start of the session. All predictors were computed in 1-second bins. For each predictor, data were pooled across animals and a linear regression model was fitted to the pooled data. The adjusted R² and p-value for the regression slope are reported. For visualization purposes only, the data was down sampled when plotting.

To calculate time constant (tau), we fitted a saturating exponential function of the form A × (1 − e^(−t/τ)) to the z-scored photometry signal beginning at the time of the first lick, where A is the plateau amplitude and τ is the time constant representing the time to reach 63.2% of the plateau. Fitting was performed using nonlinear least squares optimization.

To decompose CALCR neuron activity into slow and fast components, licking bouts were first identified as described above. The slow component was estimated by extracting z-score values during inter-bout intervals, smoothing these values with a 180-second moving average window, and interpolating across licking periods to generate a continuous slow signal across the trial. The fast component was then obtained by subtracting the slow component from the raw z-scored trace. The z-score of the slow component was defined as the mean absolute z-score across the full trial, and the z-score of the fast component as the mean absolute z-score during licking bouts only.

#### Conditioned taste avoidance to GDF-15

To validate chemogenetic silencing of GFRAL neurons, a conditioned taste aversion paradigm was used. *Gfral::hM4Di* mice were water deprived for the five-day duration of the experiment, except for two hours just prior to dark phase. All experiments were conducted during the light phase. On pairing days (days 1 and 3), mice were given 30-minute access to sucralose, after which they received an IP injection of GDF-15 paired with either vehicle or DCZ. On intervening days (days 2 and 4), mice were given 30-minute access to water. On the test day (day 5), mice were given simultaneous access to two sippers, one containing sucralose and one containing water, for 30 minutes. Licks on each sipper were recorded and preference for sucralose was expressed as the fraction of total licks directed to the sucralose sipper.

### Chemogenetic experiments

#### Behavior

All experiments were counterbalanced for stimulation order and included within-animal (saline vs DCZ) controls. *Gfral:hM4Di* experiments also included genotype (±DREADDS) controls, with genotype controls being littermates lacking the Cre or reporter allele. Animals were tested in behavioral chambers (Med Associates) without food or water access, and chambers were cleaned with 70% ethanol between sessions to remove olfactory cues from previous experiments. Before experiments began, mice underwent an overnight session with ad libitum access to water and chow, followed the next day by a second session in which animals received a saline injection and then licked Ensure in the chamber.

For Ensure, glucose 24%, and Intralipid 10% licking experiments, as well as for chow experiments, mice were given access to each compound for at least 24 hours in their home cage a few days prior to the experiment to prevent neophobia during the experiment. Prior to the experiment, mice were fasted overnight. On the experimental day, mice were injected with either deschloroclozapine (DCZ, 0.3 mg/kg, Hello Bio) or saline vehicle and then placed in the behavioral chamber for a 10 min acclimation period prior to food presentation. For water consumption experiments, animals were water-deprived overnight, and the same protocol was followed.

#### Analysis

For bout analysis, licking behavior was analyzed from raw lickometer data files. A licking bout was defined as a continuous sequence of at least 3 licks in which no inter-lick interval exceeded 5 seconds. Total lick count, bout number, and mean bout duration (average length of all lick bouts) were computed for each animal across the full 60-minute session. Cumulative lick counts were computed in 10-second bins aligned to the time of lickometer access. Cumulative lick traces were averaged across animals within each group and plotted as mean ± SEM over the 60-minute session. To measure chow consumption, the pellet was weighed before and after the experiment.

### Optogenetic experiments

#### Laser parameters

For continuous and pre-stimulation protocols, light was delivered at 20 Hz using 10-ms pulses in a 2-second on, 3-second off pattern. For closed-loop experiments during ingestion, a 2-second train of light was triggered by each detected lick, monitored using a contact lickometer interfaced with Med Associates Med-PC software. Light was delivered through a single fiber optic patch cord (Doric, MFC_200/240-0.22_6mm_ZF1.25(G)_FLT) connected to a 473-nm DPSS laser (Shanghai Laser and Optics Century BL473-100FC). Laser power was set to 10 mW, measured at the tip of the patch cord prior to each experimental session.

#### Behavior

All experiments were counterbalanced for stimulation order and included within-animal (±laser) and genotype (±opsin) controls, with genotype controls being littermates lacking the Cre or reporter allele. Animals were tested in behavioral chambers (Med Associates) without food or water access, and chambers were cleaned with 70% ethanol between sessions to remove olfactory cues from previous experiments. Before experiments began, mice underwent two habituation sessions: an overnight session with ad libitum access to water and chow, followed the next day by a second session in which animals were connected to the fiber optic patch cord. To prevent conditioned taste aversion during GFRAL stimulation, mice were given access to Ensure at least three times before the first stimulation experiment.

For Ensure licking experiments, mice were fasted overnight prior to testing. On the experimental day, animals were given a 10-minute acclimation period in the chamber before food presentation and stimulation. Food access and stimulation then occurred simultaneously for 60 minutes. For water consumption experiments, animals were water-deprived overnight, and the same protocol was followed. For pre-stimulation experiments, animals first habituated in the chamber for 10 minutes, then received continuous stimulation for 60 minutes in the absence of food. Following stimulation, animals were given 60 minutes of access to Ensure without further light delivery.

For open-loop stimulation experiments measuring chow consumption, animals were fasted overnight before the experiment. After 10 min habituation, animals received a pellet of standard chow (PicoLab 5053) for self-paced consumption over 60 min. Animals also received open-loop stimulation (described above) during the entire session or no laser treatment.

#### Analysis

Licking bout analysis was done as described above for chemogenetics. In addition, to the parameters above, latency to first lick was computed for each animal across the full 60-minute session. Cumulative lick counts were computed in 10-second bins aligned to the time of lickometer access. Cumulative lick traces were averaged across animals within each group and plotted as mean ± SEM over the 60-minute session. To measure chow consumption, the pellet was weighed before and after the experiment.

### Fluoroscopy experiments

#### Fluoroscopy setup

Fluoroscopy recordings were done using a custom-built low energy fluoroscope (Glenbrook Technologies, Newark, NJ). Fluoroscopy data was acquired at 30 frames/s with a voltage of 25 kV and a current of 0.216 mA. Mice were habituated to head-fixation and body restraint in a custom conical tube for at least five days prior to the first experiment. During habituation mice received IG infusions of water or Ensure. To minimize radiation exposure to mice, X-ray exposure was intermittent and lasted no more than 30 continuous seconds, and no more than 5 minutes on a single session.

For the IG infusions, either water or Ensure was mixed with the oral contrast barium suspension (barium sulfate oral suspension 60% w/v, Liquid E-Z-PAQUE, Bracco Diagnostics) for visualization under X-ray imaging, in ratio of 1:1. Infusions were delivered using a syringe pump (Harvard Apparatus, 70–2001) over 1 min or 5 min at an infusion rate of either 0.1 mL/min or 0.5 mL/min, respectively, for a total infusion volume of 0.5 mL. The i.g. catheter was attached to the syringe pump using plastic tubing and adapters (AAD04119, Tygon; LS20, Instech). For optogenetic fluoroscopy experiments, mice received continuous stimulation at 20 Hz using 10-ms pulses in a 2-second on, 3-second off pattern. Light was delivered through a single fiber optic patch cord (Doric, MFC_200/240-0.22_6mm_ZF1.25(G)_FLT) connected to a 473-nm DPSS laser (Shanghai Laser and Optics Century BL473-100FC). Laser power was set to 10 mW, measured at the tip of the patch cord prior to each experimental session

#### Experimental approach

For all experiments mice were fasted 3-4 hours before the experiments to allow for clearing of gastric contents. All experiments were counterbalanced for stimulation order and included within-animal controls.

For the initial GFRAL stimulation experiment, mice were placed in the custom headbar set up and attached to IG catheter and optogenetic patch cord. The laser was turned on 2 min prior to the start of the IG infusion. Mice received an infusion of barium/water over 5 min at an infusion rate of 0.1 mL/min. Time zero was defined as the start of the infusion. Fluoroscopy images were taken intermittently at several time points for 30 min. For the second GFRAL stimulation experiment (assessing if it changes stomach size), animals received an infusion of barium/Ensure over 1 min at an infusion rate of 0.5 mL/min. Ensure was given to avoid rapid gastric emptying of contents. The laser was then turned on 1 minute after the infusion was complete. Time zero was defined as the time the laser was turned on. Fluoroscopy images were taken intermittently at several time points for 10 min.

For CALCR stimulation experiments, animals were injected with either deschloroclozapine (DCZ, 0.3 mg/kg, Hello Bio) or saline vehicle and then 20 min later attached to IG catheter and placed in the fluoroscopy chamber. Mice received an infusion of barium/water over 5 min at an infusion rate of 0.1 mL/min. Time zero was defined as the start of the infusion. Fluoroscopy images were taken intermittently at several time points for 30 min

#### Analysis

Gastric size was assessed from real-time X-ray fluoroscopy videos using a custom MATLAB tool for frame-by-frame stomach area measurement. For each video, a calibration frame was first selected in which a ruler of known length was visible to establish a pixel-to-millimeter conversion factor. A time-zero frame corresponding to the start of the intragastric infusion or the optogenetic stimulation was then designated, and all subsequent measurements were expressed as time relative to this frame. For each selected frame, the user drew a freehand region of interest around the stomach. Within this region, the image was Gaussian blurred, normalized, and thresholded using Otsu’s method to generate candidate dark and bright regions. The region with the lower mean intensity was selected as the stomach mask, holes were filled, and small objects removed. The largest remaining connected component was retained and its area computed in square millimeters using the calibration factor. Each segmentation was visually inspected and either accepted or repeated until a satisfactory result was obtained. Stomach area was measured at multiple time points across the infusion period for each animal.

#### Fos experiment

Animals were habituated to intraperitoneal injections over five days prior to the experiment and fasted overnight before the experimental day. On the day of the experiment, *Calcr::hM3Dq* mice and littermate controls received an intraperitoneal injection of deschloroclozapine (DCZ, 0.3 mg/kg, Hello Bio). Two hours later, animals were transcardially perfused and brains were collected for sectioning and Fos staining and described below. Cell counts were performed manually on sections with matching anterior-posterior position in ImageJ (v2.14). The investigator was blinded to genotype of the animal.

#### Histology

Mice were transcardially perfused with heparinized PBS followed by 10% formalin. Brains were post-fixed overnight in 10% formalin at 4°C, then transferred to 30% sucrose in PBS overnight. Subsequently, brains were sectioned at 40 μm on a cryostat and mounted with DAPI Fluoromount-G (Southern Biotech), and then imaged using the Nikon Eclipse Ti2-E.

For rabies experiments, mCherry and GFP expression were amplified by immunohistochemistry. Floating sections were washed three times in PBS-T (0.1% Triton-X), blocked in 2% normal donkey serum in PBS-T for one hour at room temperature, and incubated overnight in primary antibodies in blocking solution at 4 °C (chicken anti-GFP, 1:2000, Abcam ab13970; goat anti-mCherry, 1:1000, Sicgen AB0040-200). Sections were then washed in PBS-T three times and incubated in secondary antibodies (donkey anti-chicken Alexa Fluor-488, 1:1000, Sigma-Aldrich SAB4600031; donkey anti-goat Alexa 547, 1:1000, LifeTech A11056) in blocking solution for one hour at room temperature, washed in PBS three times, mounted onto slides, and imaged using the Nikon Eclipse Ti2-E.

For Fos experiments the same protocol was used but with primary antibody rabbit anti-cFOS (Abcam ab190289, 1:1000) and secondary antibody donkey anti-rabbit Alexa 647 (Invitrogen A31573, 1:1000).

## Statistics

All data are presented as mean ± SEM. Statistical significance is indicated as follows: *p < 0.05, **p < 0.01, ***p < 0.001, ****p < 0.0001. P values for comparisons between two groups were calculated using the Wilcoxon signed-rank test for paired data or the Mann-Whitney U test for unpaired data. When multiple related outcomes were compared within the same experiment, P values were adjusted using the Benjamini-Hochberg procedure to control the false discovery rate. Sample sizes were not predetermined by statistical power analysis. Experiments were not randomized or blinded. Photometry, behavioral, and fluoroscopy data were analyzed in MATLAB.

---

**Extended Data Figure 1.**
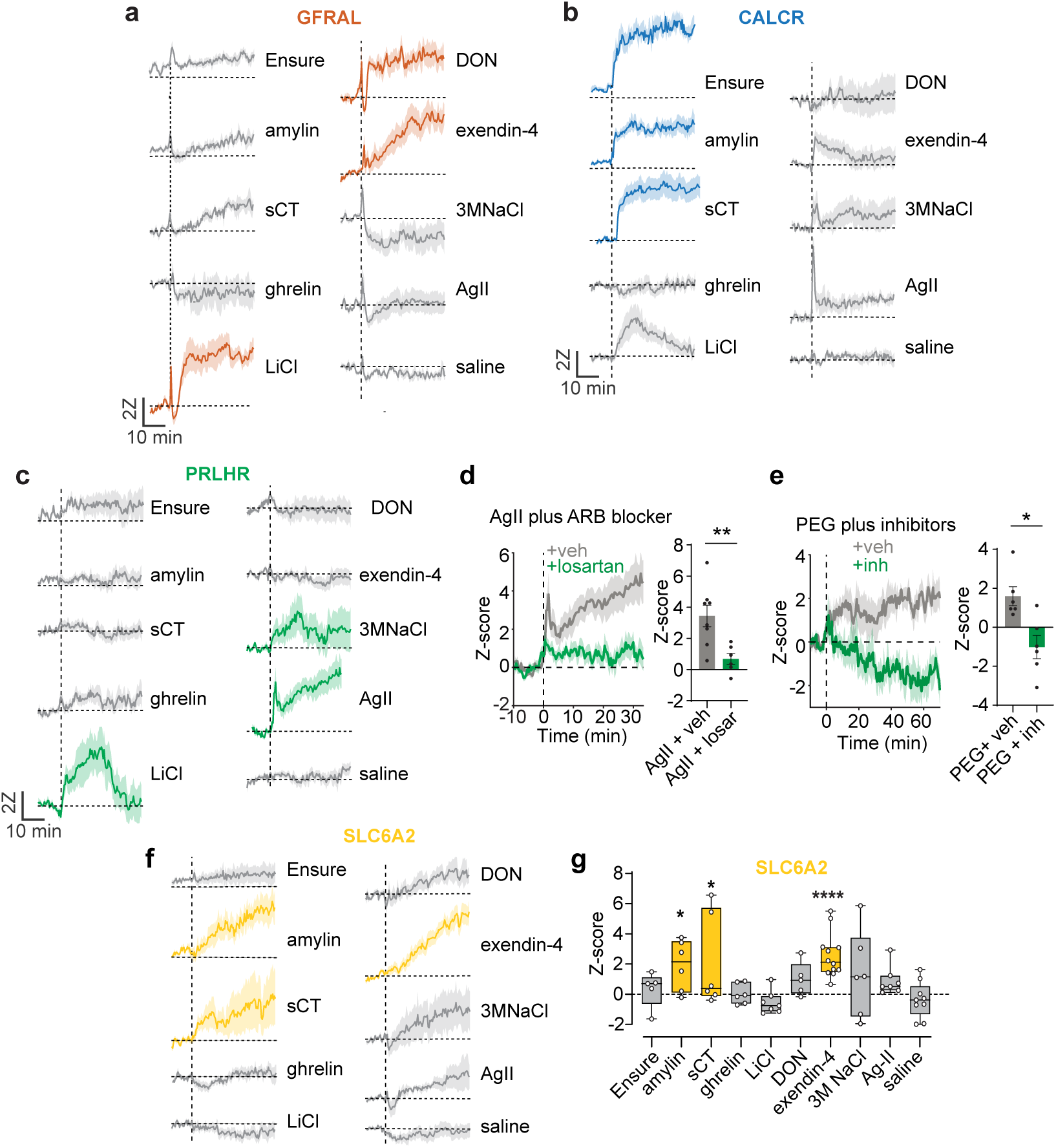
Glutamatergic cell types within the area postrema show different response profiles. **a,** GFRAL neuron response aligned to liquid food access or IP injection. Traces are mean +/- SEM. Colored traces are statistically significant responses after multiple comparison correction. **b,** CALCR neuron response aligned to liquid food access or IP injection. **c,** PRLHR neuron response aligned to liquid food access or IP injection. **d,** Left, PRLHR neuron responses to an injection of angiotensin after a subcutaneous injection of the angiotensin-receptor blocker losartan (losar) or vehicle (veh) (colors per graph on right). Right, z scores (0–40 min). **e,** Left, PRLHR neuron responses to a subcutaneous injection of the osmotic agent polyethylene glycol (PEG) which causes hypovolemia. Prior to PEG administration animals received subq losartan + IP captopril (inh) which inhibit endogenous angiotensin signaling, or vehicle (veh). Right, z scores (0–40 min). Data shows that PRLHR neurons respond to hypovolemia in an angiotensin-dependent manner. **f,** SLC6A2 neuron response aligned to liquid food access or IP injection. **g,** Boxplot showing z-score (0-40 min) for SLC6A2 neuron responses. Colored boxes are statistically significant. Statistical comparisons are relative to the baseline prior to injection/ingestion. *P<0.05, **P<0.01, ****P<0.0001

**Extended Data Figure 2.**
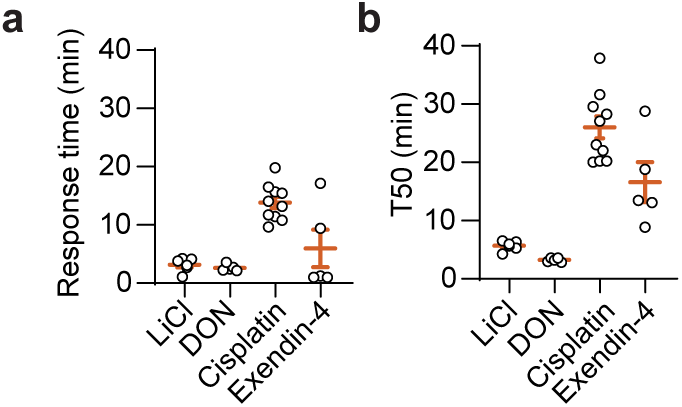
GFRAL neurons respond rapidly to emetics. **a,** Response time for various emetics tested. Response time was defined as the time at which the z-score crossed above zero and remained there for the duration of the recording (see arrows in Fig. 2b,c). **b,** Time to reach 50 percent of max z-score (T50) for different emetics tested. Data are mean ± SEM.

**Extended Data Figure 3.**
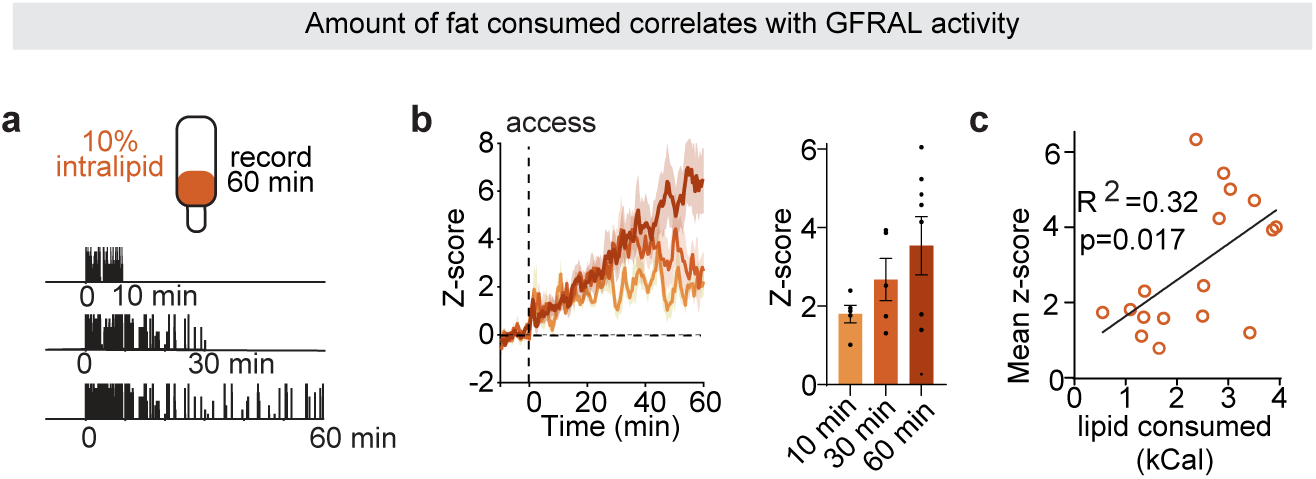
Characterization of GFRAL lipid response. **a,** Schematic, animals were given access to intralipid for 10 min, 30 min, or 60 min, and GFRAL neuron activity was recorded for 60 min. **b,** Left GFRAL neuron response after different intralipid access durations. Right, z-scores. **c,** Regression of calories of lipid consumed vs average z-score for GFRAL neurons.

**Extended Data Figure 4.**
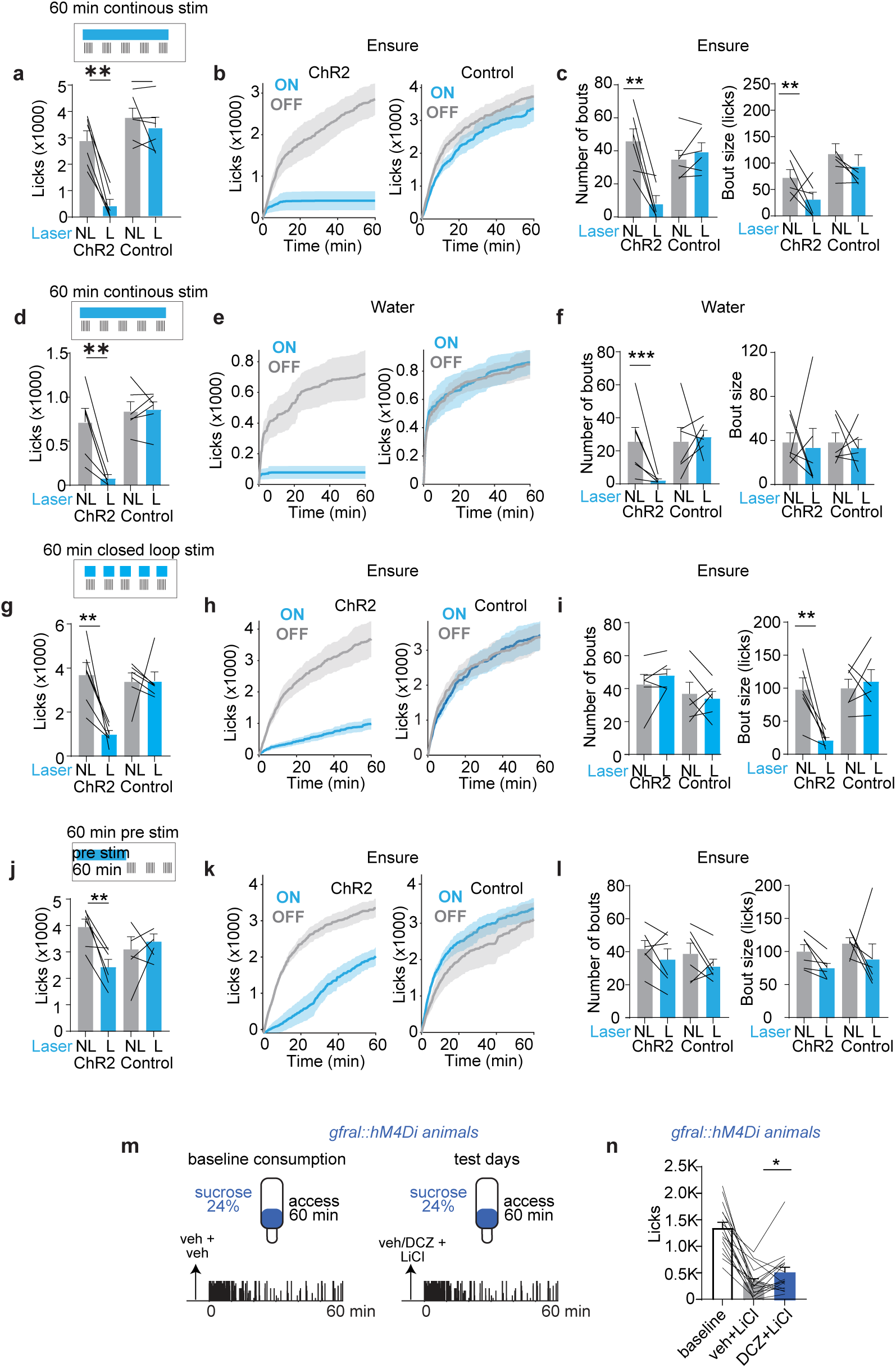
Effects of activating and silencing GFRAL neurons. **a,** Ensure consumption during continous optogenetic stimulation (60-min trial) of GFRAL neurons in either no laser (NL) or laser (L) trials for *Gfral::ChR2* and littermate controls. **b,** Cumulative licks for *Gfral::ChR2* (left) and controls (right). **c,** Number of bouts (left) and bout size (right). **d,** Water consumption during continous optogenetic stimulation (60-min trial) of GFRAL neurons in either no laser (NL) or laser (L) trials for *Gfral::ChR2* and littermate controls. **e,** Cumulative licks for *Gfral::ChR2* (left) and controls (right). **f,** Number of bouts (left) and bout size (right). **g,** Ensure consumption during closed-loop optogenetic stimulation (60-min trial) of GFRAL neurons in either no laser (NL) or laser (L) trials for *Gfral::ChR2* and littermate controls. **h,** Cumulative licks for *Gfral::ChR2* (left) and controls (right). **i,** Number of bouts (left) and bout size (right). **j,** Ensure consumption after 60 min pre-stimulation of GFRAL neurons in either no laser (NL) or laser (L) trials trials for Gfral::ChR2 and littermate controls. **k,** Cumulative licks for *Gfral::ChR2* (left) and controls (right). **l,** Number of bouts (left) and bout size (right). **m,** Experimental schematic. On the first day, *Gfral::hM4Di* animals received two injections of vehicle followed by access to sucrose. Baseline consumption was quantified. On the test days, animals received an injection of either vehicle or DCZ, followed by an injection of LiCl, and were then given access to sucrose. **n,** Number of licks in each condition. *P<0.05, **P<0.01, ***P<0.001

**Extended Data Figure 5.**
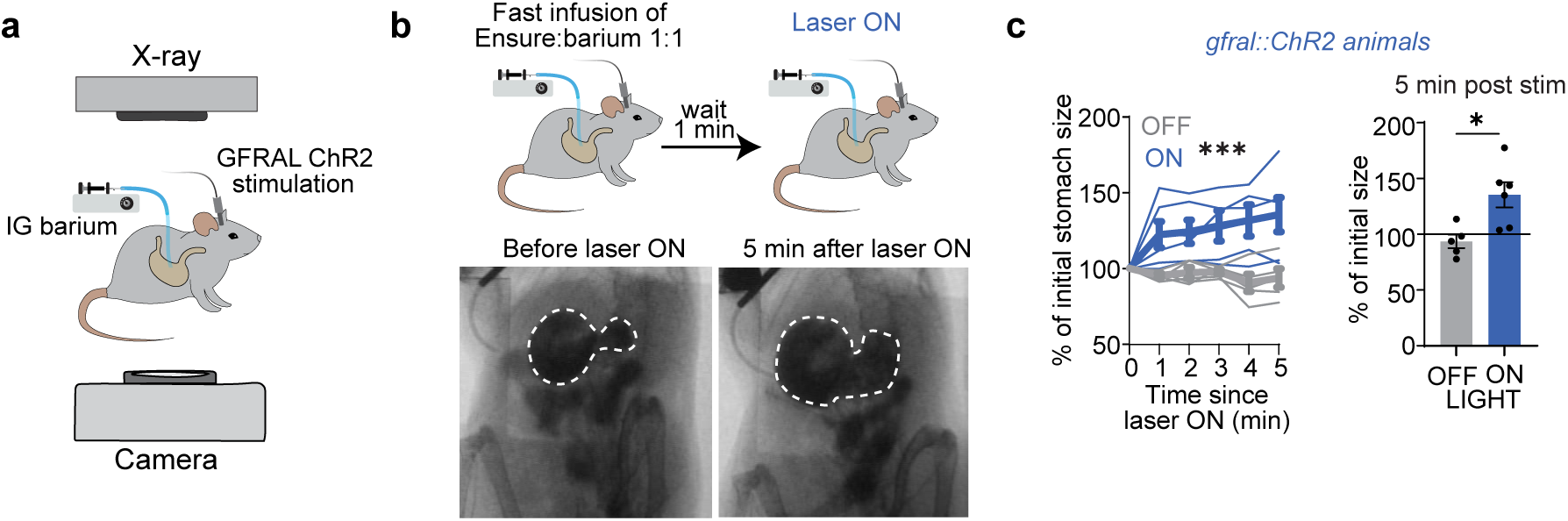
GFRAL neuron activity increase gastric size. **a,** Schematic for fluoroscopy experiments. Animals were transferred to fluoroscopy chamber and received an intragastric (IG) infusion of barium contrast. Intestinal dynamics were visualized by X-ray fluoroscopy, consisting of an X-ray source and a camera. Images were recorded at 30 Hz. **b,** Top, experimental schematic, animals received an intragastric infusion of Ensure with barium contrast (0.5 mL/min over 1 min) and 1 min later the laser was turned on to activate GFRAL neurons. Bottom, representative images of stomach (white outline) at selected time points following laser-on in *Gfral::ChR2* mice. **c,** Left, relative stomach size over time, normalized to size at the end of the infusion just prior to stimulation. Right, percentage of initial size, 5 min after stimulation. *P<0.05, ***P<0.001.

**Extended Data Figure 6.**
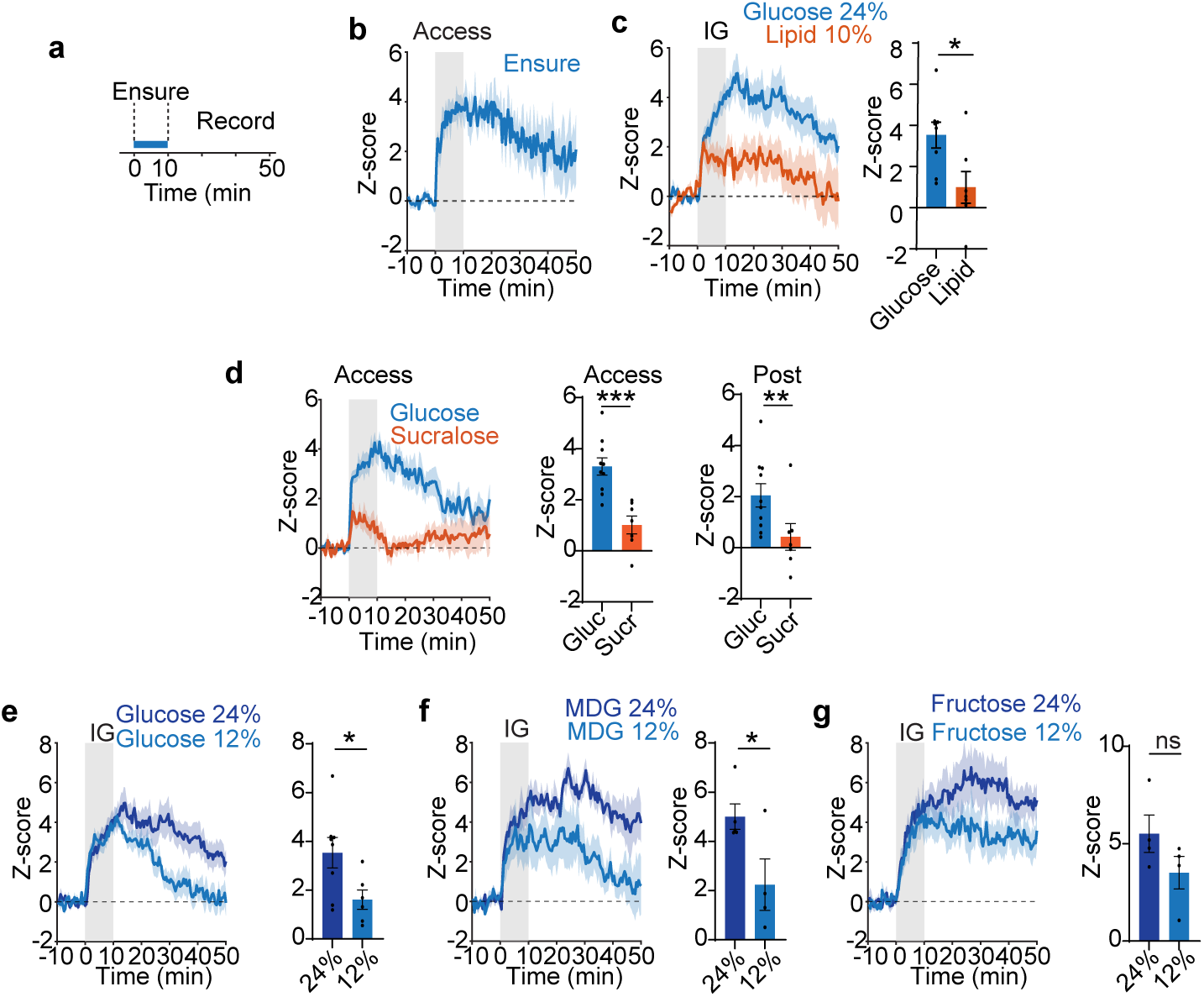
CALCR neurons show sustained responses to sugar ingestion. **a,** Schematic of intermittent Ensure access experiment. **b,** CALCR neuron activity aligned to intermittent access to liquid diet Ensure. Access was given for 10 min and then removed. **c,** CALCR neuron responses to an intragastric infusion (0-10 min, 1.0 mL) of isocaloric solutions of glucose vs. intralipid. Right, z-score (0-40 min after infusion). **d,** Left, CALCR neuron responses to 10 min access of glucose vs. non-caloric sweetener sucralose. Right, z-score during access (0-10 min) and post access (10-50 min) periods. **e,** CALCR neuron responses to an intragastric infusion (0-10 min, 1.0 mL) of glucose 24% vs. 12%. Right z-score (0-40 min after infusion). **f,** CALCR neuron responses to an intragastric infusion (0-10 min, 1.0 mL) of MDG 24% vs. 12%. Right z-score (0-40 min after infusion). **g,** CALCR neuron responses to an intragastric infusion (0-10 min, 1.0 mL) of fructose 24% vs. 12%. Right z-score (0-40 min after infusion). ns, not significant, *P < 0.05 **P<0.01, ***P<0.001. Data are mean ± SEM.

**Extended Data Figure 7.**
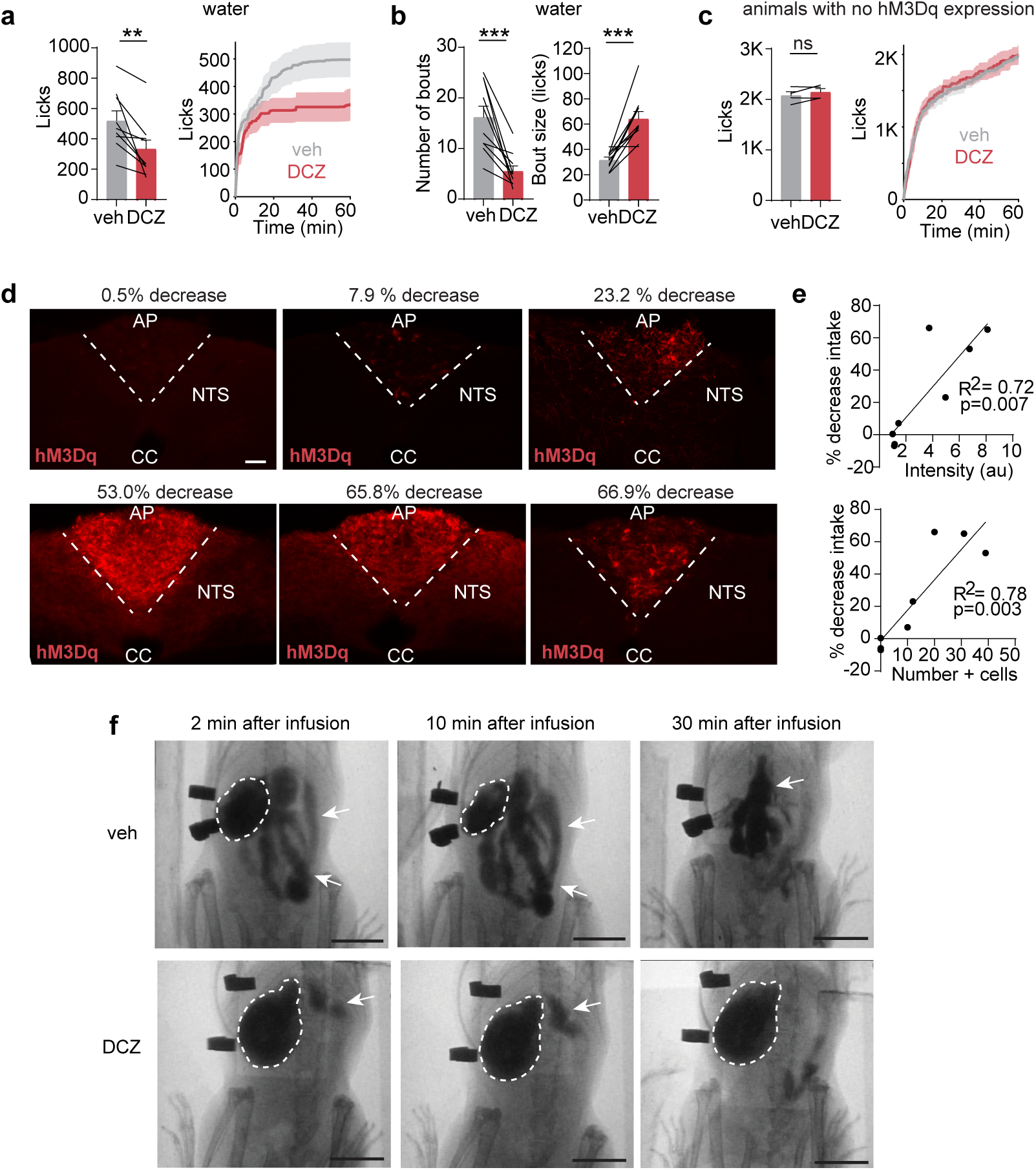
CALCR neurons pace ingestion and slow gastric emptying. **a,** Water consumption after either veh or DCZ injection in animals expressing hM3Dq::mCherry. Left, total number of licks. Right, Cumulative licks. **b,** Number of bouts and bout size during water consumption. **c,** Ensure consumption after either veh or DCZ injection in animals with no hM3Dq::mCherry expression. **d,** Varying expression levels of hM3Dq::mCherry in CALCR area postrema neurons. Number above each image indicates percentage decrease in Ensure consumption when animals are treated with DCZ compared to vehicle. Scale bar: 100um. **e,** Top. Linear regression for normalized intensity of AP vs decrease in Ensure consumption (DCZ vs saline). Bottom, Linear regression for hM3Dq::mCherry positive cells in AP vs decrease in Ensure consumption (DCZ vs saline). **f,** Representative images of stomach (white outline) and small intestines (white arrows) at different time points after infusion for animals receving either veh or DCZ prior to infusion. scale bar: 10 mm. ns, not significant, P<0.05, **P<0.01, ***P<0.001.Data are mean ± SEM.

